# Stable brain PET metabolic networks using a multiple sampling scheme

**DOI:** 10.1101/2021.03.16.435674

**Authors:** Guilherme Schu, Wagner Scheeren Brum, Yuri Elias Rodrigues, Julio Cesar de Azeredo, Tharick A. Pascoal, Andrea Lessa Benedet, Sulantha Mathotaarachchi, Pedro Rosa-Neto, Jorge Almeida, Eduardo R. Zimmer, for the Alzheimer’s Disease Neuroimaging Initiative

## Abstract

The human brain’s interregional communication is vital for its proper functioning. A promising direction for investigating how these regions communicate relies on the assumption that the brain is a complex network. In this context, images derived from positron emission tomography (PET) have been proposed as a potential source for understanding brain networks. However, such networks are often assembled via direct computation of inter-subject correlations, neglecting variabilities between subjects and jeopardizing the construction of group representative networks. Here, we used [^18^F]FDG-PET data from 1027 individuals at different syndromal stages (352 CU, 621 MCI and 234 AD) to develop a novel method for constructing stable (i.e. resilient to spurious data points) metabolic brain networks. Our multiple sampling (MS) scheme generates brain networks with higher stability when compared to the conventional approach. In addition, the proposed method is robust to imbalanced datasets and requires 50% fewer subjects to achieve stability than the conventional approach. Our method has the potential to considerably boost PET data reutilization and advance our understating of human brain network patterns in health and disease.

## 1. Introduction

Brain glucose utilization is directly proportional to synaptic activity [1]. This assumption serves as the basis for the interpretation of fluorine-18 fluoro-2-deoxyglucose positron emission tomography ([^18^F]FDG PET or FDG-PET) imaging [2]. For the past 40 years, cerebral glucose metabolism - indexed by semi-quantitative (standardized uptake value [SUV] or standardized uptake value ratio [SUVR]) or quantitative measurements - has been used to estimate regional tissue glucose uptake in normal and pathological states. More recently, derivation of metabolic brain networks (MBNs) from inter-subject [^18^F]FDG-PET [3]–[5] data has been proposed as a refined manner to gain additional information about the energetic architecture of the brain [6]–[8].

MBNs have been firstly introduced by Horwitz et al. (1984) and are typically represented by weighted graphs whose nodes are associated with predefined brain regions in which the edges reveal the network coupling (i.e. synchronicity) [9], [10]. Currently, MBNs rely on the computation of linear correlations (i.e. Pearson correlation coefficients) of [^18^F]FDG-PET measures of brain regions across subjects. This straightforward methodology has been used to identify metabolic architectural changes in schizophrenia [11], Alzheimer’s disease (AD) [12]–[15], cognitive impairment [16], [17], Parkinson’s Disease [18], [19], diabetes mellitus [20], and aging development [21], [22].

Although MBNs constructed from inter-subject correlations continues to be explored, evaluations of the variability of such networks remain neglected, leading to substantial inconsistencies evidenced across neuroimaging modalities [23], [24] and across similar [^18^F]FDG-PET studies [3], [4]. We hypothesize that inter-subjects linear group correlations are highly unstable – susceptible to outliers, sample size, and data imbalance – and may be hampering studies reproducibility. Here, we present an innovative general methodology for constructing more stable MBNs on a multiple sampling (MS) scheme in association with a more conservative method for multiple comparisons correction.

## 2. Materials and methods

### 2.1 Data source

Data used in the preparation of the present study were obtained from the Alzheimer’s Disease Neuroimaging Initiative (ADNI) database (http://adni.loni.usc.edu). The ADNI is a longitudinal study with approximately 50 sites across the United States, and Canada launched in 2003, led by Principal Investigator Michael W. Weiner, MD. The main goal of ADNI is to identify how imaging and fluid biomarkers, neuropsychological and clinical assessments can be used to understanding, predict and stage AD. This effort’s aim is to determine sensitive and specific biomarkers to aid researchers and clinicians in developing new treatments, as well as monitoring their effectiveness while reducing the expense and duration of clinical trials.

The ADNI’s research population consists of cognitively unimpaired (CU), early or late mild cognitive impairment (MCI) and Alzheimer’s clinical syndrome (AD) individuals. The follow-up duration of each group is specified in the protocols for ADNI-1, ADNI-2 and ADNI-GO study phases. The institutional review boards of all sites participating in the ADNI provided review and approval of the ADNI data collection protocol. Written informed consent was obtained from all participants at each site. For up-to-date information, see www.adni-info.org.

### 2.2 Study participants selection

We selected 1027 individuals (352 CU individuals; 621 MCI individuals and 234 AD dementia individuals) who underwent [^18^F] FDG-PET and structural MRI scans in ADNI-1, ADNI-GO and ADNI-2 phases of the ADNI project. Demographic characteristics of the groups are presented in Table 1. Additionally, in order to tune the proposed method parameters, the dataset was split into train and test sets as depicted in Table 2.

**Table 1.** Demographics.

|  | CU | MCI | AD |
| --- | --- | --- | --- |
| **Number of subjects (%)** | 352 (31.8%) | 641 (60.4%) | 234 (22.8%) |
| **Age, y, mean (SD)** | 74.26 (5.62) | 72.71 (7.51) | 74.34 (7.68) |
| **Male, no (%)** | 177 (50.28) | 380 (59.28) | 135 (57.69) |
| **Education, y, mean (SD)** | 16.28 (2.79) | 16.00 (2.76) | 15.45 (2.93) |
| **MMSE, mean (SD)** | 29.02 (1.17) | 27.77 (1.74) | 23.69 (2.38) |
| **APOE ε4 carriers, no (%)** | 97 (27.55) | 315 (49.14) | 157 (67.09) |

**Table 2.** Dataset organization.

|  | No. Individuals x No. VOIs | Group |
| --- | --- | --- |
| **Train set** | 300 X 72 | MCI |
| **Test set** | 352 X 72 | CU |
|  | 321 X 72 | MCI |
|  | 234 X 72 | AD |

### 2.3 Neuroimaging methods

MRI and PET acquisitions followed the ADNI’s protocols (http://adni.loni.usc.edu/methods). MRI T1-weighted images were preprocessed for gradient unwarping and intensity normalization [25]. The T1-weighted images were then processed using the CIVET image-processing pipeline and registered using a nine-parameter affine transformation and nonlinearly spatially normalized to the MNI 152 template [26]. [^18^F]FDG-PET images were preprocessed to have an effective point spread function of full-width at half-maximum of 8 mm. Subsequently, linear registration and nonlinear normalization to the MNI 152 template were performed with the linear and nonlinear transformation derived from the automatic PET to MRI transformation and the individual’s anatomical MRI coregistration. [^18^F]FDG-PET SUVR maps were generated using pons as the reference region [27]. Further details on our processing pipeline can be found elsewhere [27], [28].

### 2.4 Construction of MBNs using a multiple sampling scheme

Let *X* ∈ ℝ^*N*×*d*^ be a *N* × *d* dataset matrix containing PET SUVR values of *N* subjects for *d* volumes of interest (VOIs) of a given group. The proposed MS scheme consists of generating *n* samples *Y*^1^, …, *Y*^*n*^ of the dataset *X*. A general bootstrap sample *Y*^*k*^ ∈ ℝ^*N*×*d*^ is denoted as a dataset matrix containing repeated rows of the original dataset (i.e. *Y^k^* ⊆ *X*). In this context, each column of *Y*^*k*^ is denoted by the vector 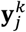 (with 1 ≤ *j* ≤ *d*).

Given the aforementioned notations, we can construct the adjacency matrix *M*^*k*^ ∈ ℝ^*d*×*d*^, associated with the dataset *Y^k^*, by computing the Pearson linear correlation coefficients between all columns of matrix *Y*^*k*^, with *p*, *q* = 1, …, *d*, as follows:

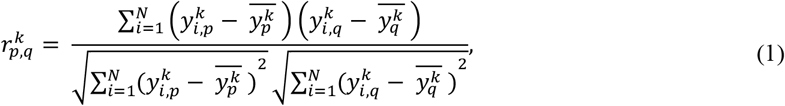

where 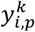 and 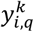 correspond to the *i* -th element of the vectors 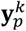 and 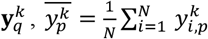 and 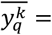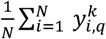 are the mean values of the column vectors 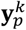 and 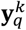, respectively.

For each dataset *Y^k^* generated by the MS scheme, we compute, using Eq. 1, a weighted undirected network 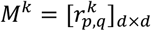. Given a collection of generated networks Φ = {*M*^1^, *M*^2^, …, *M*^*n*^} for a group of interest, we then decide which of these networks can be chosen as the representative network of the group. A direct approach is to compute the mean matrix 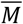 (i.e. 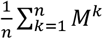) and identify which of the matrices in the set Φ best approximates 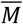 according to some metric. Hence, we define a representative network 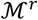 of the group as:

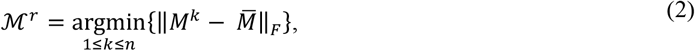

where || . ||_*F*_ is the Frobenius norm. With this formulation, we seek to find the adjacency matrix 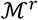 that best approximates a generated matrix *M*^*k*^ from the mean matrix 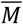.

Algorithm 1 can be used iteratively to construct group representative networks. In practice, a sample *Y*^*k*^ of the original dataset *X* can be generated using different sampling approaches, such as bootstrap and subsampling methods. Likewise, different criteria than the one presented in Eq. 2 (i.e. the mean matrix) can be used to choose the representative network of the group of interest. Additionally to the mean criterion (see Eq. 2), in Section 3, we test the median criterion and a generalization of the Bayesian mode criterion proposed in our previous work [29]. Fig. 1 summarizes the MS approach. Finally, given the collection of generated networks Φ for a group of interest, the representative MBN is chosen and corrected for multiple comparisons using false discovery rate (FDR) [30]. For more information about how to optimally set a subsampling scheme or mode criterion see supplemental materials.

**Figure 1:**
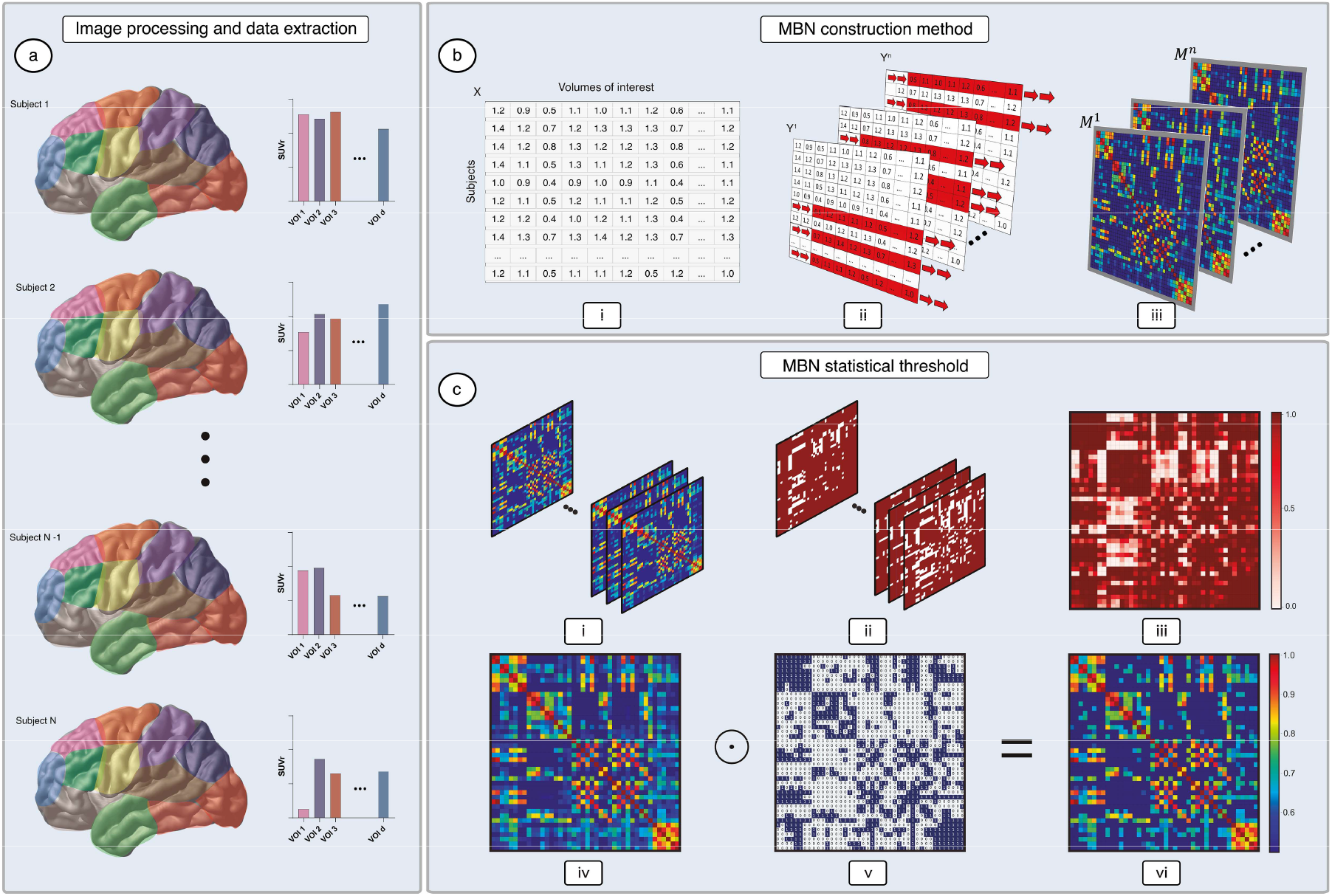
Illustration of the multiple sampling scheme. [^18^F]FDG-PET standardized uptake value ratio (SUVR) are computed for *d* Volumes of interest (VOIs) and *N* subjects - as shown in (a) – generating a dataset matrix for a group of interest. The original dataset matrix *X* ∈ ℝ^*N*×*d*^ shown in (b.i) is bootstrap sampled *n* times, generating the datasets *Y*^1^, …, *Y*^*n*^ shown in (b.ii). For each of the sampled datasets *Y*^1^, …, *Y*^*n*^, adjacency matrices *M*^1^, …, *M*^*n*^ are constructed by computing inter subject Pearson correlation coefficients (b.iii). The collection of FDR corrected adjacency matrices *M*^1^, …, *M*^*n*^ shown in (c.i), have their entries evaluated using Eq. 3, generating the binary matrices shown in (c.ii). An average degree probability distribution matrix 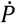 (i.e. a probability map), shown in (c.iii), is computed by averaging the collection of binary matrices. Given a defined threshold *θ*, matrix 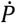 is then thresholded using Eq. 4, generating a threshold matrix 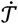 shown in (c.v). Finally, the FDR corrected representative matrix 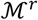 shown in (c.iv) is thresholded by computing the Hadamard product between 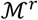 and 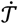, which results in the corrected matrix shown in (c.vi).

**Algorithm 1.**
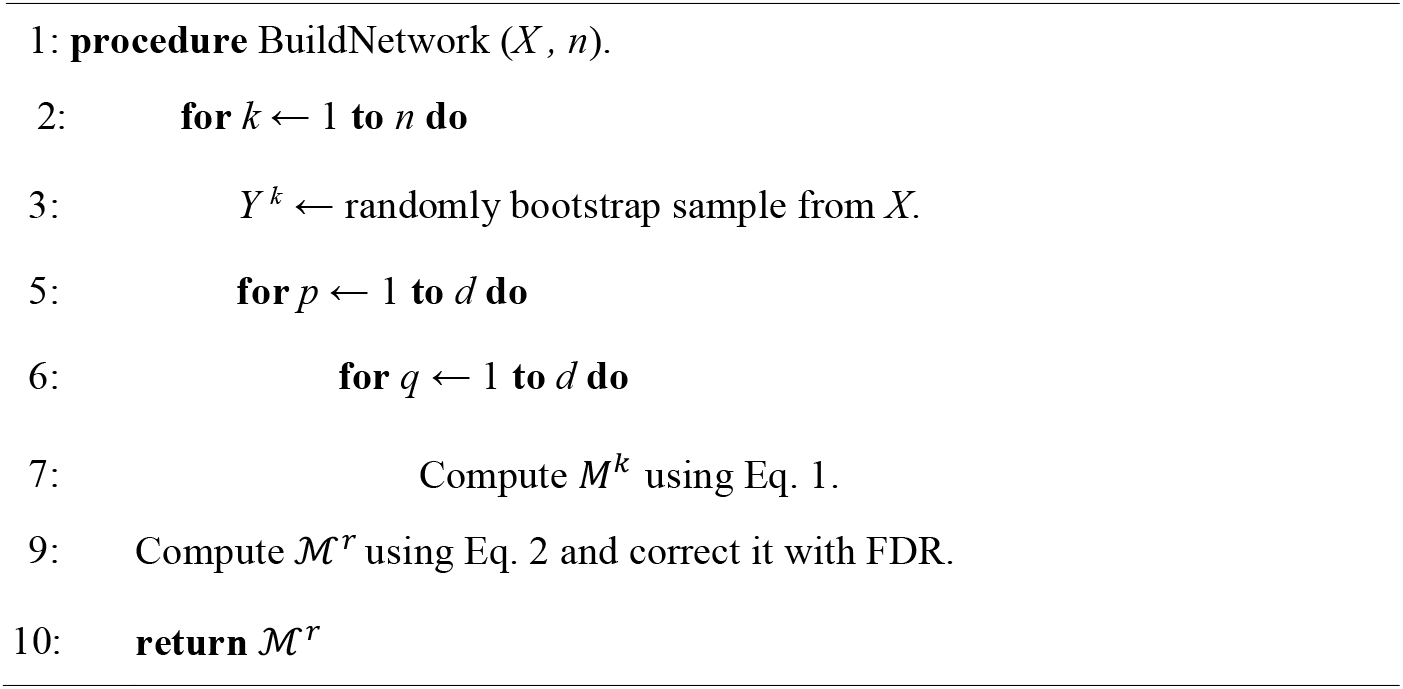
Metabolic brain network construction

### 2.5 Network threshold using probability maps

Here, we propose a novel approach for thresholding the group representative MBNs in addition to the FDR method. We generate a probability map (Pmap) based on the average degree distribution computed from *n* bootstrap samples. We verify which edges are more likely to occur after sampling the data randomly multiple times. With this scheme, if the probability of an edge to occur is higher than a threshold *θ*, one should maintain that edge, otherwise, one should disregard it. Explicitly, for *p*, *q* = 1, …, *d*, we define the entries of the probability matrix 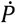 as:

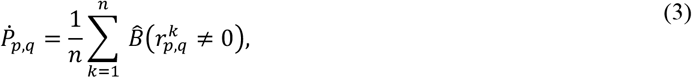

where 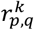 corresponds to elements of *M*^*k*^ (corrected by FDR) and the function 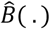 evaluated for an arbitrary logical statement z given by:

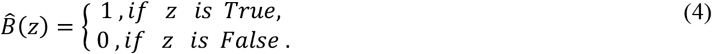

Given a choice for *θ*, we generate a threshold matrix using the computed probability map as:

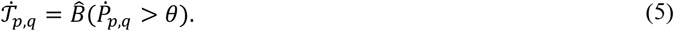

Finally, we threshold the FDR corrected representative matrix 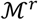 by computing the Hadamard product with the threshold matrix 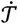 (i.e. 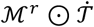). In practice, we define *θ* = 1 − *α*, where *α* is the statistical threshold typically used in multiple comparison methods such as FDR. Fig. 1c shows a representation of the proposed threshold scheme. Using the collection of networks generated with the MS approach (shown in Fig. 1c.i), we test whether their entries are different than zero, generating the binary matrices showed in Fig. 1c.ii. Computing the sum over all binary matrices and dividing their elements by *n*, we generate the probability matrix 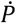 (shown in Fig. 1c.iii). Finally, the representative matrix corrected by FDR 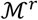 (shown in Fig. 1c.iv) is thresholded by computing the Hadamard product between 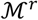 and 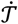 (Fig. 1c.v), which results in the corrected matrix (shown in Fig. 1c.vi).

### 2.6 Network model outlier attack

Here, we adapted the node attack strategy used in [31], and proposed an attack method that is driven by the inclusion of outliers to a group dataset aiming to evaluate MBNs stability. More specifically, we generate *Q* perturbed versions 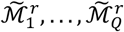 of the original representative matrix 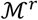. In this model, for each perturbed network, we introduce *L* outliers to the dataset *Y*^*r*^ (that has produced 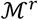) to generate a perturbed network 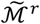. We defined outliers as data points randomly sampled from datasets belonging to the groups being compared, which are introduced in the dataset of a group of interest. In this model, if one would make an attack to the CU group, for instance, it would consist of randomly sampling data points from the MCI and AD groups to later introduce these data points in the CU dataset. In our experiments, the attacks were programmed in a way that given the parameter percentage of outliers (*P_o_*), half of the outliers introduced to the dataset of interest (e.g. CU) are from one group (e.g. MCI), and half are from the other group (e.g. AD). We understand this is a valid attack model because the metabolic patterns of different groups usually do not differ greatly in absolute value, and for that reason, might not be detectable in typical outlier-removal methods. In the experimental setup we have defined *P_o_* ∈ [2%, 5%, 8%] and the total number of attacks equal to 256.

Likewise, it is possible to apply this procedure to generate perturbed inter-subject correlation matrices with no random samples (what we refer here as the conventional method). In this case, the same outliers used to build perturbed representative matrices constructed using the MS scheme are included in the original dataset *X* ∈ ℝ^*N*×*d*^, generating a perturbed dataset 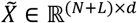. An inter-subject correlation matrix and its perturbed version can then be obtained by computing the Pearson linear correlation coefficients using *X* and 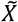 followed by FDR correction.

### 2.7 Network stability measures

We address the problem of measuring the stability of a MBN by evaluating the similarity between an arbitrary group representative network 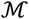 and its perturbed version 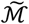. We argue that alterations in the topology and alterations in the characteristics of the network, provoked by spurious data samples, are directly associated with the stability of the network. To quantitatively evaluate the stability of a network and its perturbed version, we explore four different metrics: Hausdorff distance, Frobenius norm, Euclidean distance and Canberra distance.

The Frobenius norm can be computed directly from the network and its perturbed version (i.e. 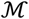 and 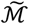). This norm is a common measure used to compute matrix similarities, with lower values (i.e. closer to 0) indicating a higher degree of network stability (i.e. matrix similarity). Likewise, we investigate the use of the Hausdorff distance to estimate network stability. Given an adjacency matrix 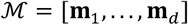 and a perturbed version of it 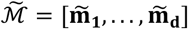, the Hausdorff distance between these two matrices is defined as [33]:

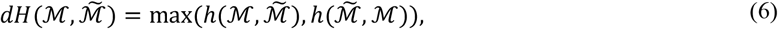

where the directional Hausdorff distance *h*( . ) is defined as:

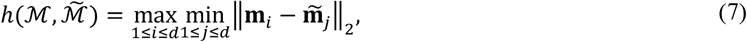

and || . ||_2_ is the Euclidean norm.

In addition, we also investigate the Euclidean and the Canberra distances as measures of network stability. In this scheme, the similarity of adjacency matrices is evaluated in the vector space spanned by the network graph theoretical measures. For each pair of networks 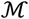, and its perturbed version 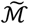 we compute feature vectors **f** = [*ge*, *ac*, *ad*, *as*, *d*, *acc*] and 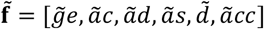, respectively. The feature vector entries correspond to graph theoretical measures known as the global efficiency (*ge*), assortativity coefficient (*ac*), average degree (*ad*), average strength (*as*), density (*d*), average clustering coefficient (*acc*). For a complete description of how to carry out their computation see further reference [34]. Given the aforementioned definitions, one can evaluate the stability of a network in terms of feature vectors with the Euclidean norm:

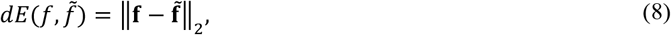

and in terms of the Canberra distance:

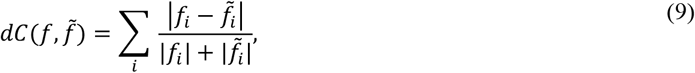

where | . | denotes the *L*_1_ norm and, *f_i_* and 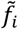 correspond to the *i*-th elements of vectors **f** and 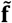, respectively. The lower the values (i.e. closer to 0) for *dE* and *dC* the higher the similarity between features, indicating a higher degree of network stability [35].

### 2.8 Data imbalance and number of subjects

Group differences in the sample size (i.e. imbalance distribution of data) are common when studying the AD spectrum. Not rarely, the number of subjects diagnosed as belonging to the MCI group is greater than the number of subjects belonging to the CU or ADs groups in the publicly available databases.

As described in Section 2.3, after constructing a representative MBN for a group of interest with the MS scheme, we correct the adjacency matrix with the FDR and the probability map (Pmap) methods, which depend on the threshold *α*. Since the number of samples tends to yield small p-values (i.e. p-value ≪ *α*) [36], we investigate, as well, the effects of data imbalance in the construction of MBNs by randomly undersampling and oversampling the [^18^F]FDG-PET data in our experiments. The random undersampling technique consists in removing instances from the larger groups until achieving the same size between all groups [37]. On the contrast, oversampling techniques typically generate synthetic patterns based on the vicinity of the existing data points, being carried until the smaller groups approximate the largest group’s size. In this work, we have used the adaptive synthetic sampling approach for imbalanced learning (ADASYN) [38] method in our experiments.

Likewise, the construction of more reproducible MBNs tends to be very dependent on the total number of subjects in a dataset. A great number of samples (i.e. subjects) produce better estimates of the underlying distribution of the data, which in turn can be used to construct more reliable and reproducible MBNs. In our experiments, we also evaluate the effects of the number of subjects as a function of the stability of the MBNs. For these experiments, we perturbed the MBNs 256 times, set *P_o_* ∈ [2%, 5%, 8%] and let *n* ∈ [10, 15, 20, …, 200].

### 2.9 Parameter tunning

As previously described in Section 2.4, the proposed MS bootstrap MBN construction scheme requires as input the number of samples (*n*), which defines the number of different networks that will be constructed to estimate later the representative matrix of a given group of interest. Aiming at optimizing our method, we assumed that the parameter *n* was independent of any group, which entails that optimizations carried out in the space of parameters of one group (e.g. MCI) could be applied to the other groups (e.g. CU and AD) with no great losses in performance. The adopted optimization scheme, iteratively computed the Bhattacharyya distance (dB) between the degree distribution (i.e. the normalized histogram of edges connecting paired nodes) of the networks generated with the MS bootstrap scheme when *n* = *k*, against the degree distribution of MBNs generated when *n* = *k* + 100 (with *k* ∈ [100, 200, …, 9900]). We optimized our setup by choosing the parameter *n* that minimized the computed Bhattacharyya distance values.

## 3. Results

### 3.1 Asymptotic decay of MBN degree distribution variability

Using the training set, our experiments have revealed that the Bhattacharyya distance values are minimized when *n* = 9300. Using the test set, we also investigated the similarity of degree distributions and weights distribution (i.e. the normalized histogram of weights connecting paired nodes divided into 100 bins) for all groups. As *n* → ∞ the degree distribution as well as the weights distribution variability, asymptotically approximates zero for all groups (Fig. 2a,b). This decay pattern was observed for the MS subsampling scheme as well (see Supplementary Fig. 1). In addition, the AD group requires larger *n* values to present dB measures comparable to CU and MCI groups when analyzing MBNs degree distribution. Moreover, we noted that the optimized parameter *n* obtained for the training set, also produced consistently low dB values (i.e. close to zero) for the CU, MCI and AD groups in the test set. Fig. 2c-e shows the stable MBNs constructed with optimized MS scheme for CU, MCI and AD groups, respectively. A list with all VOIs used to construct the MBNs can be found in Supplementary Table 1.

**Figure 2.**
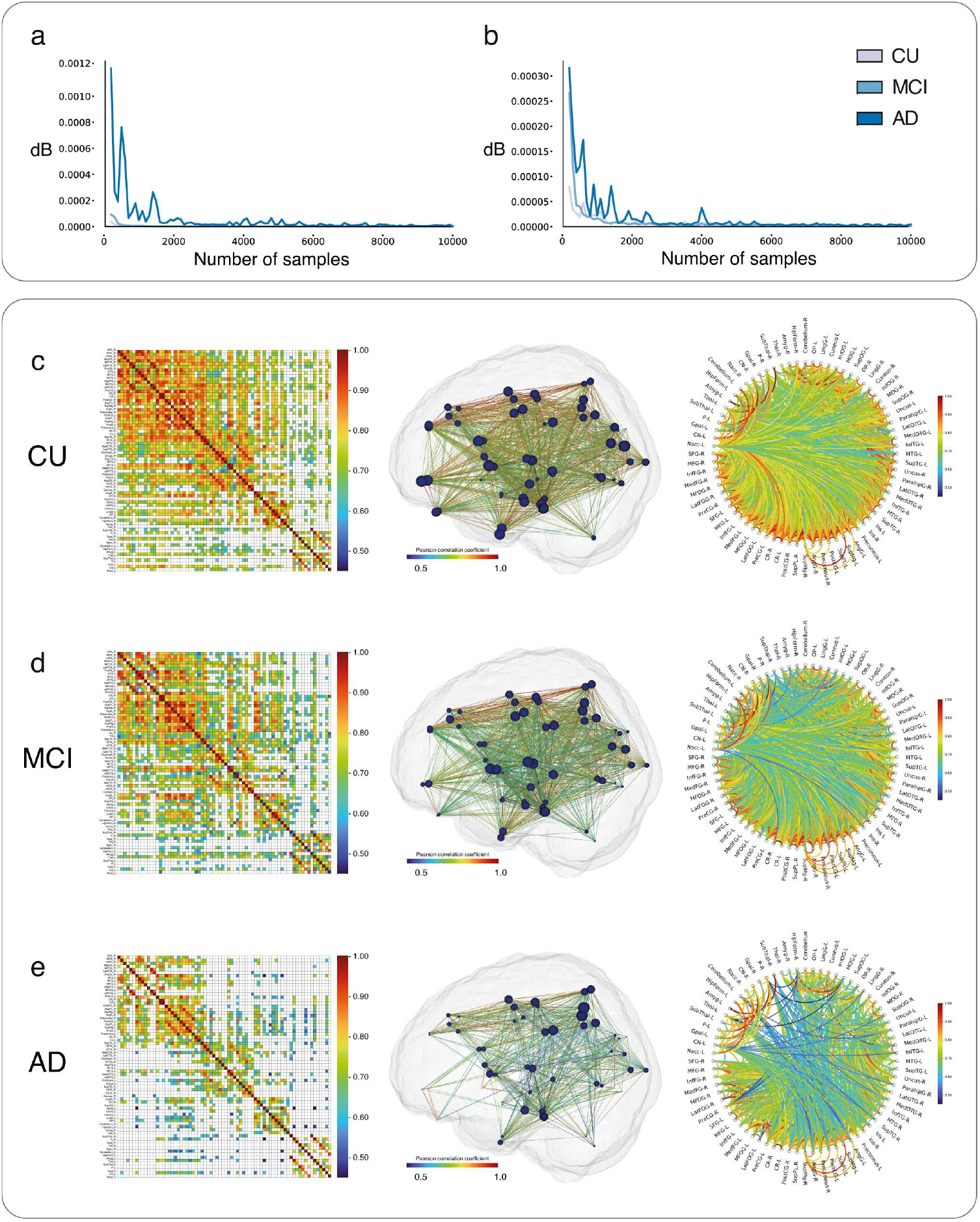
Multiple sampling bootstrap scheme assembled metabolic brain networks with optimal parameters setup. In order to define the optimal number of samples *n*, dB values were computed between distributions generated with the MS bootstrap scheme when *n* = *k*, against distributions computed when *n* = *k* + 100 (with *k* ∈ [100, 200, …, 9900]). dB distribution values for CU, MCI and AD mean representative MBNs degree distributions (a) and weights distributions (b) are displayed as a function of the number of samples. Adjacency matrices of correlation coefficients between brain regions, 3D brain surfaces displaying metabolic brain network (MBNs) architecture and circle plot visualizations are shown for CU (c), MCI (d) and AD (e) groups.

### 3.2 Probability map threshold excludes MBNs weakly linked edges

The probability map threshold (FDR + Pmap) revealed to be more conservative (i.e. it tends to eliminate more edges connecting pair of nodes) when compared with the FDR method alone. As *α* decreases, the proposed threshold tends to maintain only edges strongly linked – edges with weights values closer to |1| – for all tested groups. We also identified that the MBNs global graph measures are also impacted by different *α* values. More specially, lower *α* values led to decreased density and global efficient (GE) measures but increased assortativity (AC) and average clustering coefficients (ACC). Variations in the characteristics of the MBNs when comparing FDR+Pmap against FDR can be observed directly by inspecting the adjacency matrices of groups CU (Fig. 3a,b), MCI (Fig. 3d,e) and AD (Fig. 3g,h) as a function of parameter *θ*.

**Figure 3.**
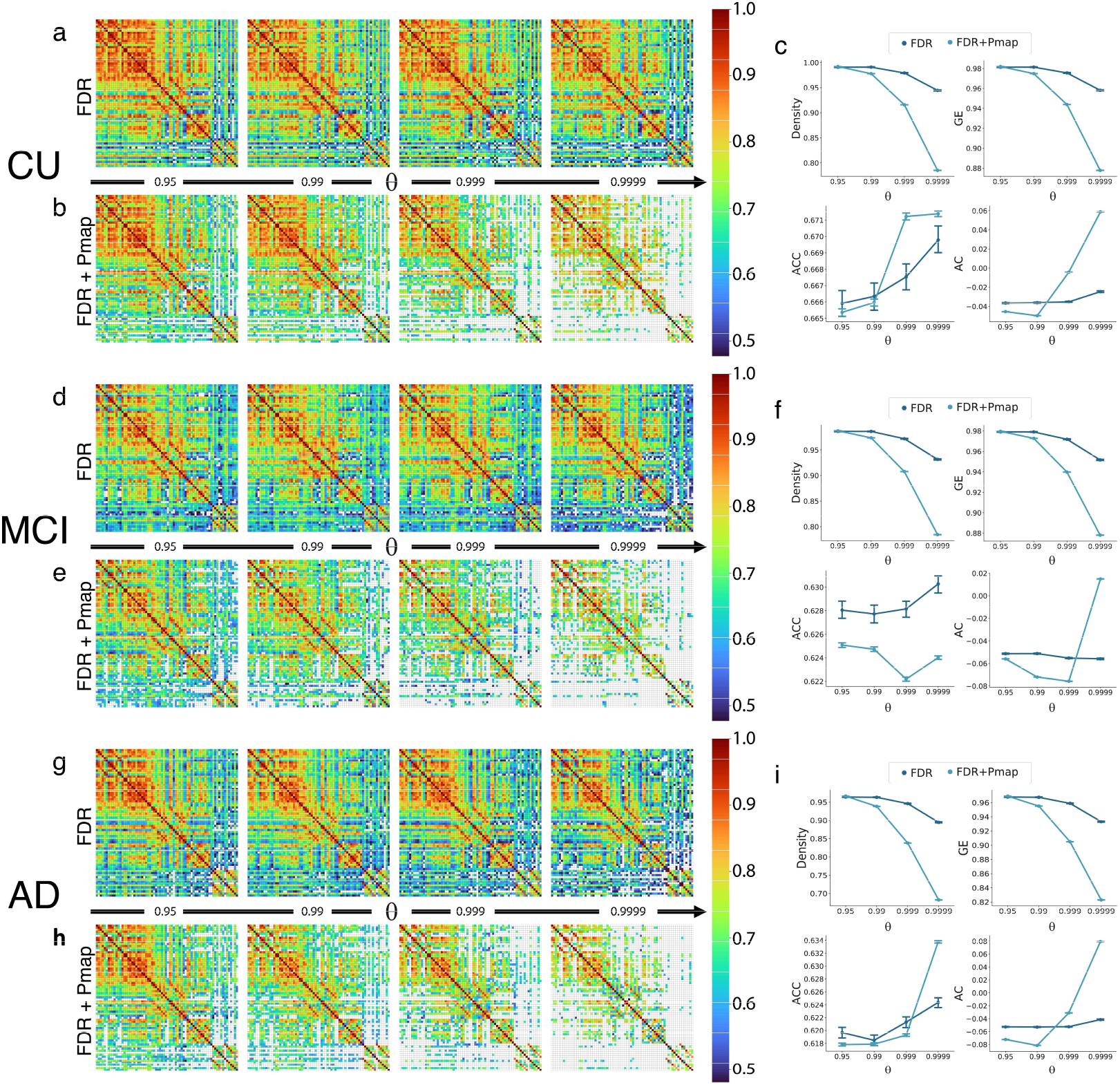
Illustration of the effects of applying the proposed thresholding method in the mean representative MBNs of groups CU, MCI and AD. (a) and (b) show, respectively, the mean representative MBN for the CU group corrected by FDR and by the FDR + Pmap approach at significance levels (alpha) 0.05, 0.01, 0.001 and 0.0001. Similarly, (d) and (e) present, respectively, the mean representative MBN for the MCI group and corrected by FDR and by the FDR + Pmap and (g) and (h) show, respectively, the mean representative MBN for the AD group corrected by FDR and by the FDR + Pmap approach. (c), (f) and (i) show the overall behavior of the graph measures Density, Global Efficiency (GE), assortativity coefficient (AC) and average clustering coefficient (ACC) as a function of the significance levels 0.05, 0.01, 0.001 and 0.0001, for the CU, MCI and AD groups, respectively.

### 3.3 MS scheme does not depend on the MBN construction criteria

In Section 2.3, we have defined the representative MBN for a group of interest as the one that best approximates the mean matrix using the Frobenius norm. We also explored other reasonable choices for the group representative matrix, namely: the mode (maximum density) matrix and the median matrix for groups CU, MCI and AD (Fig. 4a-c). Statistical analysis revealed that the mode, mean and median criteria do not differ from each other in terms of stability measures dE, dF, dH and dC (Fig. 4d-g). We computed 4 two-way analysis of variance (ANOVA), one for each stability measure, with a group factor (with 3 levels: CU, MCI and AD) and criteria factor (with 3 levels: mode, mean and median), followed by Bonferroni correction. No significant alterations were found varying the MBN construction criteria for dE (F_(2,2295)_ = 0.1639, p = 0.8488), dF (F_(2,2295)_ = 1.079, p = 0.3403), dH (F_(2,2295)_ = 1.143, p = 0.3189) and dC (F_(2,2295)_ = 0.2085, p = 0.8118). In contrast, the group factor significantly modified the stability measures dE (F_(2,2295)_ = 23.09, p < 0.0001), dF (F_(2,2295)_ = 135.1, p < 0.0001), dH (F_(2,2295)_ = 101.1, p < 0.0001) and dC (F_(2,2295)_ = 10.36, p < 0.0001) (as shown in Fig. 4d-g).

**Figure 4.**
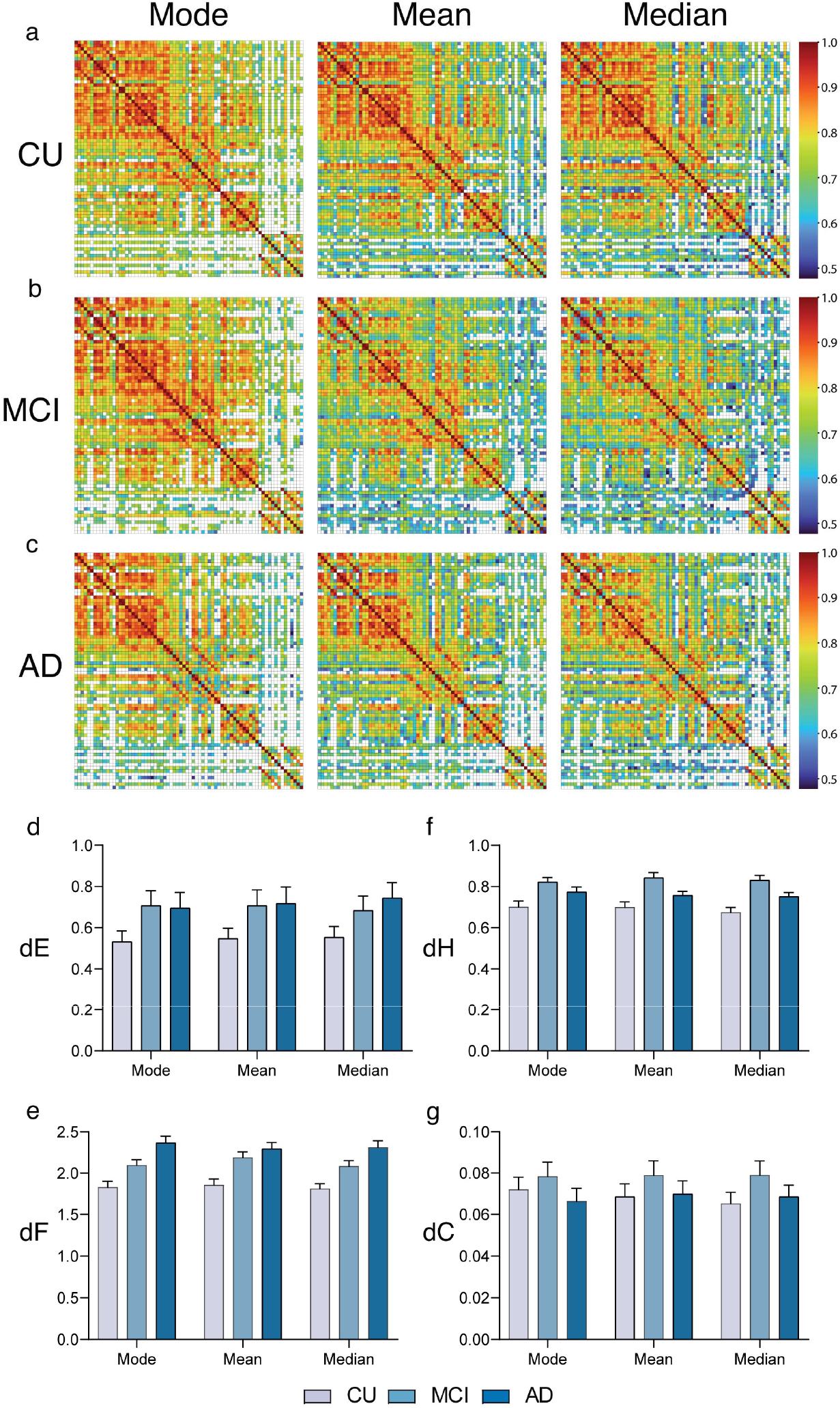
Group MBNs constructed with the mode, mean and median criteria. Representative matrices using the three construction criteria for CU, MCI and AD groups are shown, respectively, in (a), (b) and (c). CU, MCI and AD MBNs stability evaluation using the Euclidean distance (dE), the Frobenius distance (dF), the Hausdorff distance (dH) and the Camberra distance (dC) for the mode, mean and median construction criteria, are shown in (d), (e), (f) and (g), respectively.

### 3.4 Stable MBNs are robust to the data imbalance problem

Network outlier attacks (with percentage of outliers of 2%) revealed that MBNs constructed with the MS bootstrap scheme are more stable than conventional MBNs in different contexts of group imbalance (Fig. 5a-d). We performed 12 two-way ANOVAs, one for each stability measure (dE, dF, dH, dC) and one for each data balance setup (ADASYN, Undersampled, Imbalanced), followed by Bonferroni correction. For each ANOVA, the sources of variations considered were group (with 3 levels: CU, MCI and AD) and the MBN construction method (with 2 levels: conventional and bootstrap). All 12 two-way ANOVAs revealed that the construction method altered the stability measures significantly (p < 0.0001). Likewise, group and the interaction group x method were found to alter the stability measures dE, dH, dF and dC significantly (p < 0.0001). Full ANOVA results can be found in Supplementary Table 2. Likewise, the MS bootstrap scheme revealed to be more stable against network outlier attacks with 5% and 8% of outliers (Supplementary Figs. 2,3). Similarly, the MS subsambling scheme revealed to be more stable than the conventional method against network outlier attacks with 2%, 5% and 8% of outliers (Supplementary Figs. 4-6).

**Figure 5.**
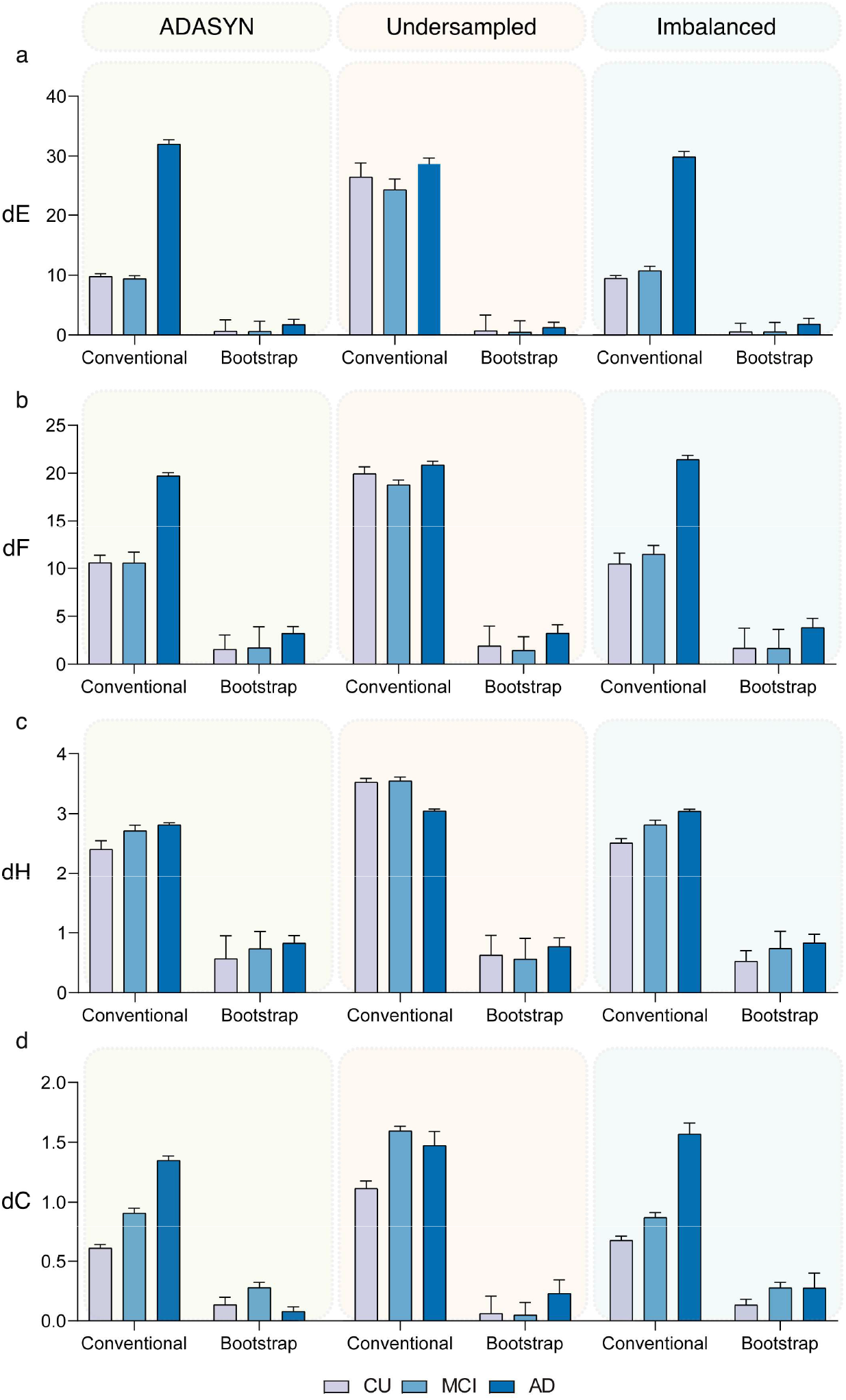
MBNs stability comparison of conventional and the MS bootstrap method for different balance schemes. Groups CU, MCI and AD were network outlier attacked 256 times (Po = 2%) with balance schemes: ADASYN (left column), Undersampled (central column) and Imbalanced (right column). MS bootstrap MBNs were constructed using the mean matrix criterim, alpha = 0.0001 and n was set to 9300. MBNs stability evaluation using the dE, dF, dH and dC, for both conventional and bootstrap methods, are shown, respectively, in (a), (b), (c) and (d).

### 3.5 Construction of stable MBNs with small datasets

Evaluation of MBN stability (with 2% of outliers attacks) as a function of the dataset size (i.e. the number of individuals used for constructing MBNs) revealed that the MS bootstrap scheme generated greater stability for groups composed by a lower number of individuals (Fig. 6a-d). More specifically, the conventional method requires in general a greater number of samples (between 100 and 200 samples) to provide a stable network, whereas the MS scheme needs fewer samples (between 50 and 100 samples) to reach stability (Fig. 6a-d). In summary, average network stability measures (dE, dF, dH and dC) computed for the MS bootstrap scheme converged faster to their minimum stability values than the conventional construction method. Likewise, the MS bootstrap scheme revealed to be more stable against network outlier attacks with 5% and 8% of outliers as a function of the dataset size (Supplementary Figs. 7,8). Similarly, the MS subsambling scheme revealed to be more stable than the conventional method against network outlier attacks with 2%, 5% and 8% of outliers (Supplementary Figs. 9-11) as a function of the dataset size.

**Figure 6.**
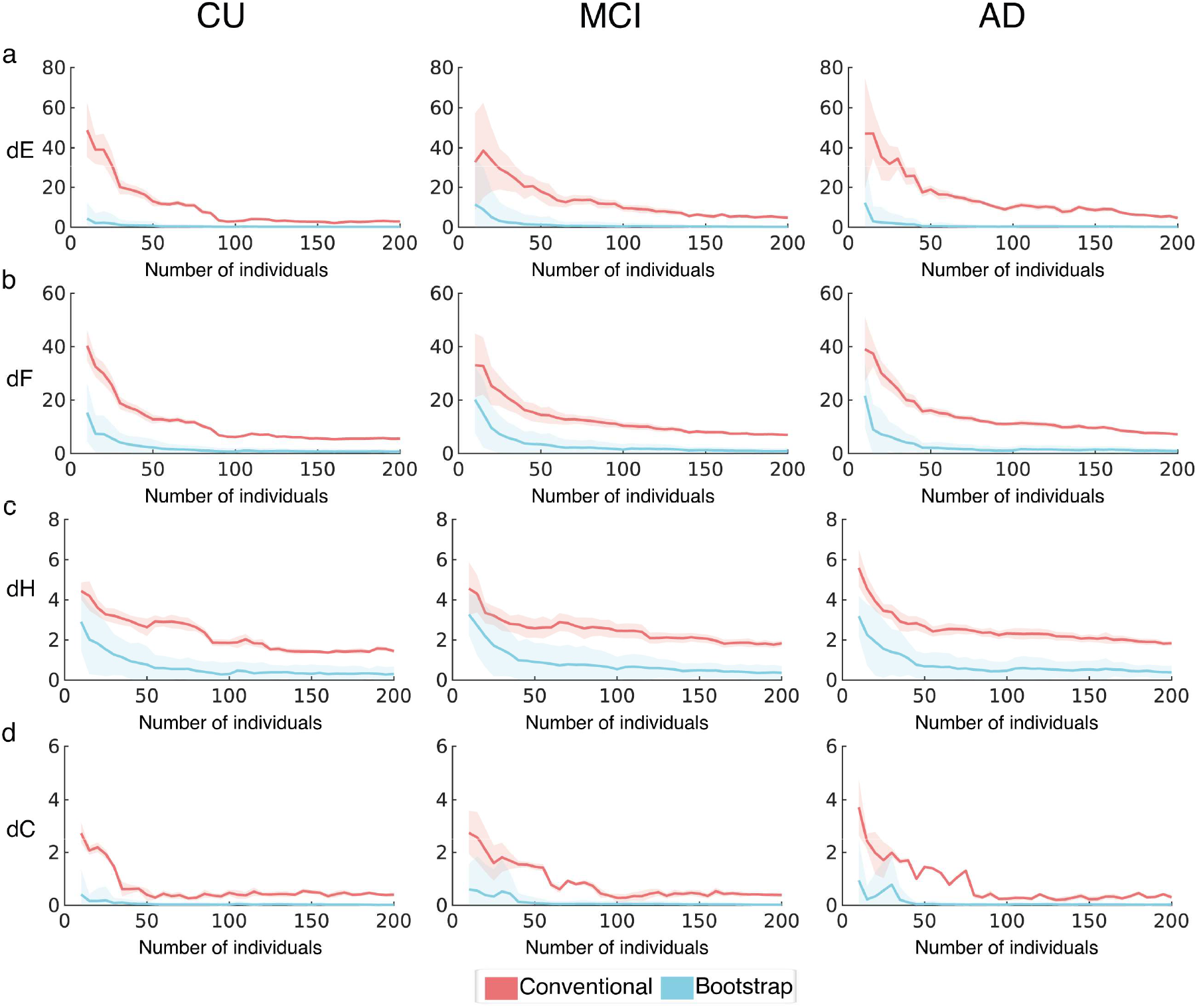
MBNs stability measures comparison of conventional and MS bootstrap method as a function of the dataset size (i.e. number of individuals). Groups CU, MCI and AD were network outlier attacked 256 times (Po = 2%) for each dataset size defined in the interval [10, 15, 20, …, 200]. MS bootstrap MBNs were constructed using the mean matrix criterion, alpha = 0.05 and n was set to 9300. MBNs stability evaluation using the dE, dF, dH and dC as a function of the number of individuals, for both conventional and bootstrap methods, are shown respectively in (a), (b), (c) and (d). Bold lines correspond to mean stability measure values and plotted light shadows correspond to 3 times the standard deviation from the mean measure.

## 4. Discussion

This report proposed a novel MS scheme to construct stable MBNs indexed by an innovative stability strategy. We showed that by sampling a dataset of [^18^F]FDG-PET images it was possible to automatically select, among many assembled MBNs, a group representative MBN that tends to preserve its characteristics and its overall organization. Additionally, the generated MBN was more resilient to outlier attacks. We also have introduced a novel thresholding method that uses the intersubject variability of the degree distribution of the assembled MBNs to maintain only connecting edges that are highly probable to exist among subjects of a given group of interest.

It is often difficult to evaluate MBNs stability since the ground truth is not known. Thus, defining a stability criterion of an MBN is an ill-posed task. In this report, we avoided the problem of defining a unique MBN ground truth for CU, MCI, and AD groups by extending the idea of node attack [31] and defining an outlier attack model to perturb a group of [^18^F]FDG-PET data, similarly to the approach proposed in [32]. The adopted strategy was based on previous work [40], and it is in agreement with a general understanding of the concept of differential privacy [41], which relies on the idea that few data points samples should not alter much the output of an algorithm. Using the framework proposed by Soundarajan and colleagues, we addressed the problem of measuring the MBN’s stability by evaluating the similarity between a group representative network and its correspondent perturbed version [35]. We tested our method using traditional similarity metrics in the field of pattern recognition. More specifically, to compare the general similarity between adjacency matrices, we have used the Frobenius norm and the Hausdorff distance. In addition, to compare the characteristics of the MBNs, we extracted graph theoretical measures, and compared the representative MBN and its perturbed version in the feature space spanned by their graph measures using the Euclidean and Canberra distances as measures of similarity, with small values indicating greater MBN stability.

Notably, we have not found statistically significant differences in the stability measures between the mean, mode and median MBN construction criteria. This indicates that the assembled MBNs may be Gaussian distributed, forming a cluster in a high dimensional manifold. Under this view, identifying the group representative MBN is equivalent to find the centroid of this Gaussian shaped cluster, which explains why there are no significant differences in the stability measures between these criteria. It is worth noting that the proposed MS scheme does not rely specifically on the connectivity measure of choice. Here, for simplicity, we have used the Pearson correlation measure – which is widely used for constructing MBNs in brain PET imaging studies – to introduce the overall formalism of this approach. Nevertheless, the MS scheme could be easily adapted to embrace other types of measures currently being tested by the brain PET imaging community, such as partial and Spearman correlations, as well as measures more sensible to nonlinear interactions such as mutual information and the maximal information coefficient [42]. Likewise, the adopted stability evaluation strategy can be used independently of how the adjacent matrices, i.e. the MBNs, may be organized for different choices of the connectivity measure. In this report, we did not fully explore the vast domain of available similarity measures between matrices in the literature. The stability results obtained with the MS scheme (see Fig. 5 and Fig. 6) revealed a great potential for assembling stable MBNs, i.e., MBNs assembled with the proposed scheme were significantly more stable than the conventional method. Additional similarity measures were not tested, but based on our findings, they will likely behave as the metrics evaluated by us.

Obtained results also have shown that the MS scheme tends to be robust to the data imbalance problem, revealing low variability between groups CU, MCI and AD for different scenarios regarding group imbalance (ADASYN, Undersampled and Imbalanced). Indeed, the overall dispersion of each group is smaller across imbalanced scenarios for the bootstrap MS scheme when compared to the conventional method. One could argue that the MS scheme may be a proper method for studying imbalanced datasets from large-scale clinical trials, such as the ADNI study. Furthermore, our findings shed light on the potential of bootstrap MS scheme to reduce the number of individuals needed to assemble group representative MBNs with improved stability, which could make MBNs a much more common secondary outcome in brain PET studies. In addition, it is worth noting that the MS scheme is application-independent. It could be used to study different brain diseases using quantitative or qualitative measures and it offers the possibility to assemble brain networks derived from multiple PET radiotracers. In addition, since the proposed scheme does not rely on data dependent parameters, it can potentially be used in other mammalian species such as non-human primates and rodents.

In summary, in the present study we provided a straightforward method to assemble reliable MBNs with multiple applications in brain PET imaging research. Our method has the potential to considerably increase PET data reutilization and advance our understating of network dynamics in normal people and brain disorders.

## 5. Data and code availability

All human data used in this study was downloaded online at the Alzheimer’s Disease Neuroimaging Initiative database (adni.loni.usc.edu). Metabolic data used to generate the MBNs and conduct the experiments here presented may be available from the corresponding author upon request. An up to date implementation of the MS scheme will be available soon in an open repository.

## Acknowledgments

Data collection and sharing for this project was funded by the Alzheimer's Disease Neuroimaging Initiative (ADNI) (National Institutes of Health Grant U01 AG024904) and DOD ADNI (Department of Defense award number W81XWH-12-2-0012). ADNI is funded by the National Institute on Aging, the National Institute of Biomedical Imaging and Bioengineering, and through generous contributions from the following: AbbVie, Alzheimer’s Association; Alzheimer’s Drug Discovery Foundation; Araclon Biotech; BioClinica, Inc.; Biogen; Bristol-Myers Squibb Company; CereSpir, Inc.; Cogstate; Eisai Inc.; Elan Pharmaceuticals, Inc.; Eli Lilly and Company; EuroImmun; F. Hoffmann-La Roche Ltd and its affiliated company Genentech, Inc.; Fujirebio; GE Healthcare; IXICO Ltd.; Janssen Alzheimer Immunotherapy Research & Development, LLC.; Johnson & Johnson Pharmaceutical Research & Development LLC.; Lumosity; Lundbeck; Merck & Co., Inc.; Meso Scale Diagnostics, LLC.; NeuroRx Research; Neurotrack Technologies; Novartis Pharmaceuticals Corporation; Pfizer Inc.; Piramal Imaging; Servier; Takeda Pharmaceutical Company; and Transition Therapeutics. The Canadian Institutes of Health Research is providing funds to support ADNI clinical sites in Canada. Private sector contributions are facilitated by the Foundation for the National Institutes of Health (www.fnih.org). The grantee organization is the Northern California Institute for Research and Education, and the study is coordinated by the Alzheimer’s Therapeutic Research Institute at the University of Southern California. ADNI data are disseminated by the Laboratory for Neuro Imaging at the University of Southern California.

