## Supplemental materials for "Stable brain PET metabolic networks using a multiple sampling scheme"

**Mode group representative matrix computation**

Previously we have used the mean matrix as the representative MBN of a group of interest. As an alternative, one could use the median or the mode criterion to decide which network should be classified as the group representative one. In order to compute the mode criterion, we have generalized the optimal Bayesian approach introduced by our previous work [29]. In this method, the representative network of the group is defined as the one located at the mode of the underlying distribution of the data. We assume that the posterior probability $p(M^{k,r}|\Phi)$ of network $M^k$ to correspond to the group representative network $M^r$ (i.e. $M^{k,r}$), given the set $\Phi$, follows a mixture of matrix normal distributions :

$$p(M^{k,r}|\Phi) = \frac{\sum_{t=1}^{n} \mathcal{MN}\left(M^k|\,\bar{M}=M^t,U,V\right)\Pi_t}{\sum_{k=1}^{n}\sum_{t=1}^{n} \mathcal{MN}\left(M^k|\,\bar{M}=M^t,U,V\right)\Pi_t}, \tag{S1}$$

where $\bar{M} \in \mathbb{R}^{d\times d}$ is the mean, $U \in \mathbb{R}^{d\times d}$ and $V \in \mathbb{R}^{d\times d}$ correspond to the variances among-rows and among-columns, respectively, and the weighting component $\Pi_t$ corresponds to the frequency of occurrence of matrix $M^k$ in the underlying distribution spanned by the set $\Phi$, such that $0 \leqslant \Pi_t \leqslant 1$ and $\sum_{t=1}^{n} \Pi_t = 1$.

In the mixture, each matrix normal distribution $\mathcal{MN}(M^k|M = M^t, U, V)$ is evaluated at matrix $M^k$ and can be written as the equivalent multivariate Gaussian:

$$\mathcal{N}(\mathbf{vec}(M^k)|\mathbf{vec}(M^t), U \otimes V = \mathbb{I}) = \frac{1}{(2\pi)^{d^2/2}} \exp[-\tfrac{1}{2}(\mathbf{vec}(M^k - M^t))^T \mathbf{vec}(M^k - M^t)], \tag{S2}$$

where $\mathbf{vec}(.)$ is the vectorization operation, $U \otimes V$ is the Kronecker product of matrices $U$ and $V$, which we assume is the identity matrix $\mathbb{I} \in \mathbb{R}^{d^2\times d^2}$, and the superscript $T$ denotes the transpose operation. Using equation equation S2, we can rewrite the posterior probability $p(M^{k,r}|\Phi)$ as:

$$p(M^{k,r}|\Phi) = \frac{\sum_{t=1}^{n} \mathcal{N}\left(\mathbf{vec}(M^k)\middle|\mathbf{vec}(M^t), \mathbb{I}\right)\Pi_t}{\sum_{k=1}^{n}\sum_{t=1}^{n} \mathcal{N}\left(\mathbf{vec}(M^k)\middle|\mathbf{vec}(M^t), \mathbb{I}\right)\Pi_t}. \tag{S3}$$

In equation S2, we estimate the density at vector $\mathbf{vec}(M^k)$ by computing a linear combination of symmetric shaped Gaussian distributions (with covariance $\mathbb{I}$) centered at each mean vector $\mathbf{vec}(M^t)$. For a more detailed discussion regarding the advantages of using the above formalism see [29]. We find the representative matrix $\mathcal{M}^{k,r}$ using the Maximum a *Posteriori* (*MAP*) approach as follows:

$$\mathcal{M}^{k,r} = \underset{1\leq k\leq n}{\arg\max}\{p(M^{k,r}|\Phi)\}. \tag{5}$$

With this formulation, we seek to find the representative network $\mathcal{M}^{k,r}$, that maximizes the posterior probabilities $p(M^{k,r}|\Phi)$, and corresponds to the mode of the underlying distribution spanned by $\Phi$. For a given $M^k$, one can compute the posterior probability $p(M^{k,r}|\Phi)$ of $M^k$ being a representative matrix $M^{k,r}$ using equation S3. Once the group representative network $\mathcal{M}^{k,r}$ is determined, the network is corrected for multiple comparisons using false discovery rate (FDR).

**Construction of MBNs using multiple subsampling**

Alternatively to using bootstrap sampling, we have also tested the use of a subsampling approach. The adotpted formalism of the subsampling scheme is slightely different to the bootstrap scheme and it is described as follows: let $X \in \mathbb{R}^{N \times d}$ be the original dataset matrix containing PET measures (e.g. SUVr) of $N$ subjects (rows), for $d$ volumes of interest (columns). The multiple subsampling method consists of generating $n$ subsamples $Y^1, \ldots, Y^n$ of $X$. We denote a general subsample as $Y^k \subseteq X$ (with $1 \leq k \leq n$), where $Y^k \in \mathbb{R}^{N(k) \times d}$ and $N(k)$ is the number of subjects used to generate that matrix. In this context, we denote each column of $Y^k$ as a vector $\mathbf{y}_j^k$ (with $1 \leq j \leq d$).

Given the aforementioned notations, we can construct the adjacency matrix $M^k \in \mathbb{R}^{d \times d}$, associated with the dataset $X^{(k)}$, by computing the Pearson correlation coefficient (for $p, q = 1, \ldots, d$) as follows:

$$r_{p,q}^k = \frac{\sum_{i=1}^{N(k)} \left(y_{i,p}^k - \overline{y_p^k}\right)\left(y_{i,q}^k - \overline{y_q^k}\right)}{\sqrt{\sum_{i=1}^{N(k)} \left(y_{i,p}^k - \overline{y_p^k}\right)^2} \sqrt{\sum_{i=1}^{N(k)} \left(y_{i,q}^k - \overline{y_q^k}\right)^2}}, \quad (1)$$

where $y_{i,p}^k$ and $y_{i,q}^k$ correspond to the $i$-th element of the vectors $\mathbf{y}_p^k$ and $\mathbf{y}_q^k$, $\overline{y_p^k} = \frac{1}{N(k)} \sum_{i=1}^{N(k)} y_{i,p}^k$ and $\overline{y_q^k} = \frac{1}{N(k)} \sum_{i=1}^{N(k)} y_{i,q}^k$ are the mean values of the vectors $\mathbf{y}_p^k$ and $\mathbf{y}_q^k$, respectively.

In practice, the subset $Y^k$ of the original dataset $X$ is obtained using a random sample percentage $S$. For each subsampling, we generate randomly a $S$ value between the interval of a maximum percentage ($S_{max}$) and minimum percentage ($S_{min}$). Hence, the construction of MBNs using the multiple subsampling scheme firstly generates the percentage $S$ of subjects that will be removed from the original dataset $X$ (i.e. rows of matrix $X$) and after, determines which subjects will be randomly drawn from $X$ to generate $Y^k$. The input parameters, $S_{max}$, $S_{min}$ and the total amount of random subsamplings $n$ were evaluated using the train set and are described in the next section.

**Parameter tuning with a subsampling scheme**

MBN construction using the multiple subsampling scheme relies on the choice for the parameters of the maximum percentage ($S_{max}$) and the minimum percentage ($S_{min}$), which are used to generate randomly a sample percentage $S$. Also, our proposed construction method requires as input the number of subsamples $n$ which defines the number of different networks that will be constructed to later estimate the representative matrix of a given group of interest. In our search for optimizing these parameters, we have used the train set as described in Section 2.2. Once again, we assumed that the method's parameters were independent of any group, which means that optimizations carried out in the space of parameters of one group (e.g. MCI) could be applied to the other groups (e.g. CN and AD) with no great losses in performance.

In our experiments, we have setup $S_{max}$ ∈ [10%, 12%, …, 30%] and have fixed $S_{min}$= 0.5%. The adopted optimization scheme, at first, fixes $S_{max}$, and iteratively computes the Bhattacharyya distance between the normalized degree distribution (i.e. the normalized histogram of edges connecting paired nodes) of the networks generated with the multiple subsampling scheme with parameters $n$ = k and $n$ = k +100 (with k ∈ [100, 200, …, 9900]. When all possible values of $n$ are searched, then $S_{max}$ is incremented and the procedure is repeated. Finally, the parameters that minimize the Bhattacharyya distance value, among all, are the optimal parameters to be chosen. In our experiments we found the optimal parameters $S_{max} = 10\%$ and $n = 9300$.

### Supplemental Figures

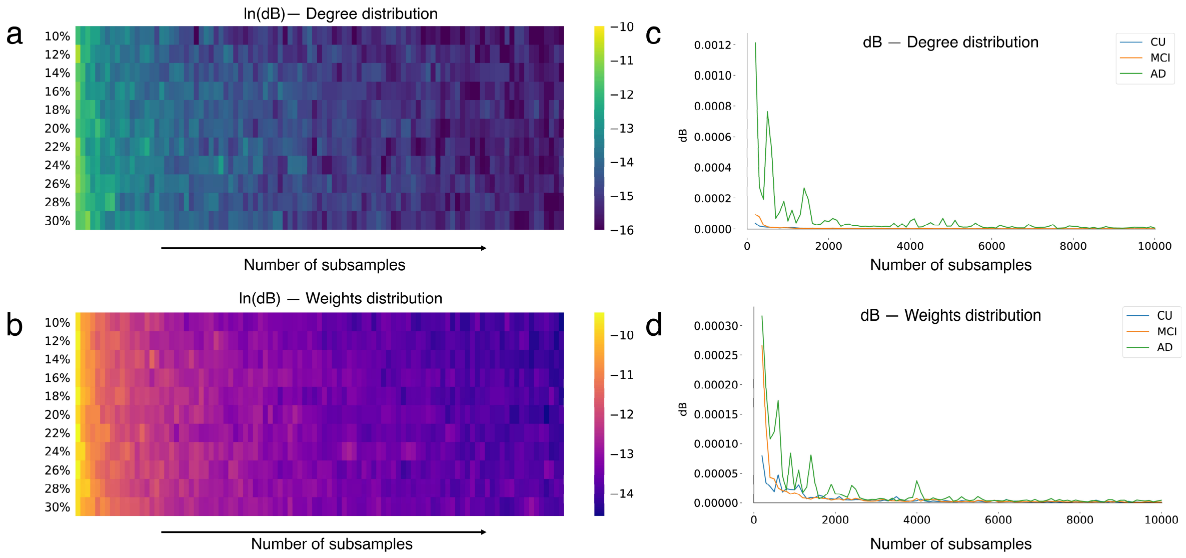


**Supplemental Figure 1. Behavior of the Bhattacharrya distance (dB) as a function of the number of samples.** Natural logarithm of dB values computed for MCI (i.e. the training set) mean representative MBNs degree distributions (a) and weights distributions (b) as a function of the number of subsamples and $S_{max}$. (c) and (d) show the behavior of the dB values computed for CU, MCI and AD (in the test set) mean representative MBNs degree distribution and weigths distributions, respectively, as a function of the number of subsamples. Setup: $S_{max} = 10\%$.

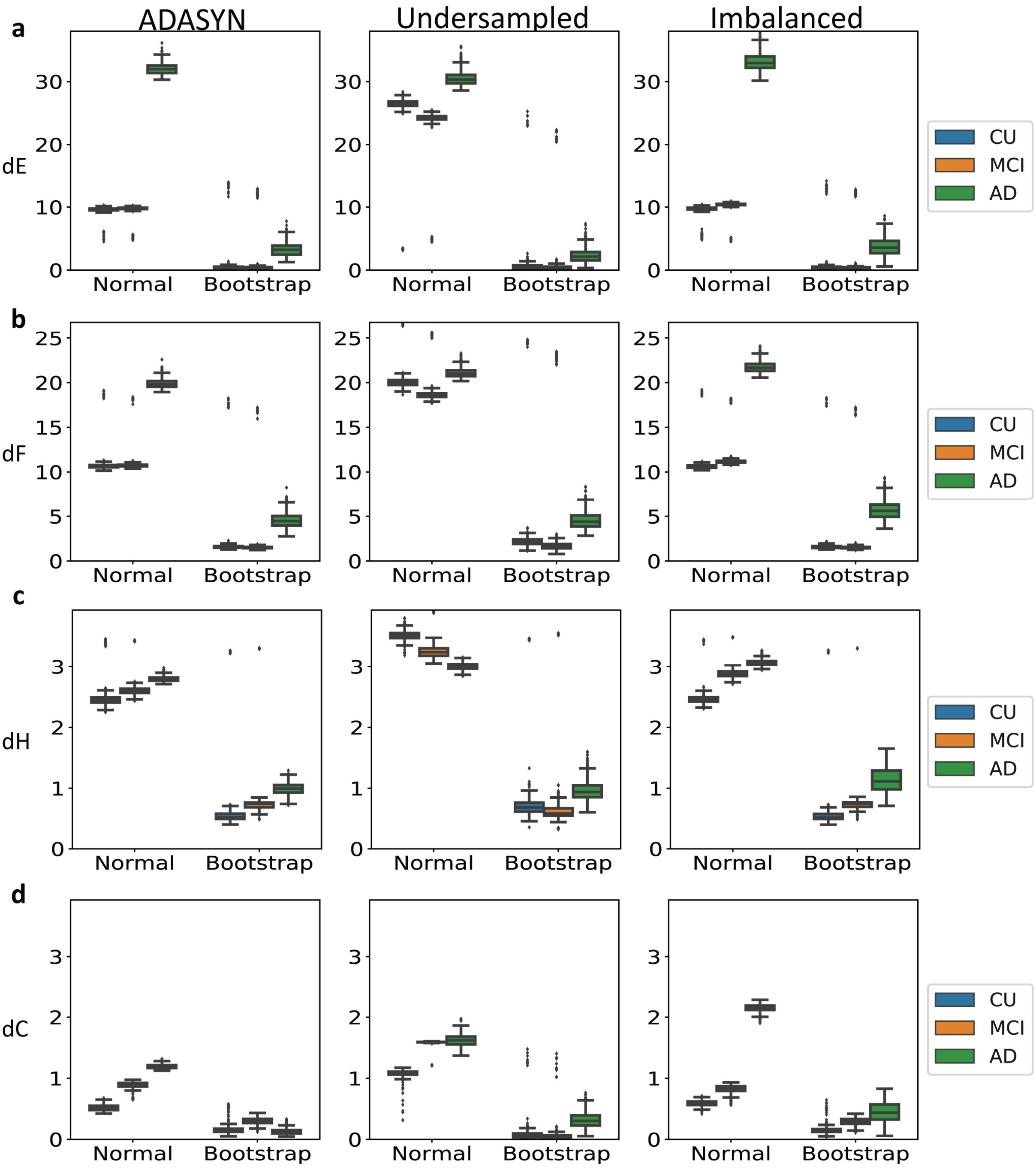


**Supplemental Figure 2. MBNs stability comparison of conventional and the MS bootstrap method for different balance schemes.** Groups CU, MCI and AD were network outlier attacked 256 times (Po = 5%) with balance schemes: ADASYN (left column), Undersampled (central column) and Imbalanced (right column). MS bootstrap MBNs were constructed using the mean matrix criterion, alpha = 0.0001 and n was set to 9300. MBNs stability evaluation using the dE, dF, dH and dC, for both conventional and bootstrap methods, are shown, respectively, in (a), (b), (c) and (d).

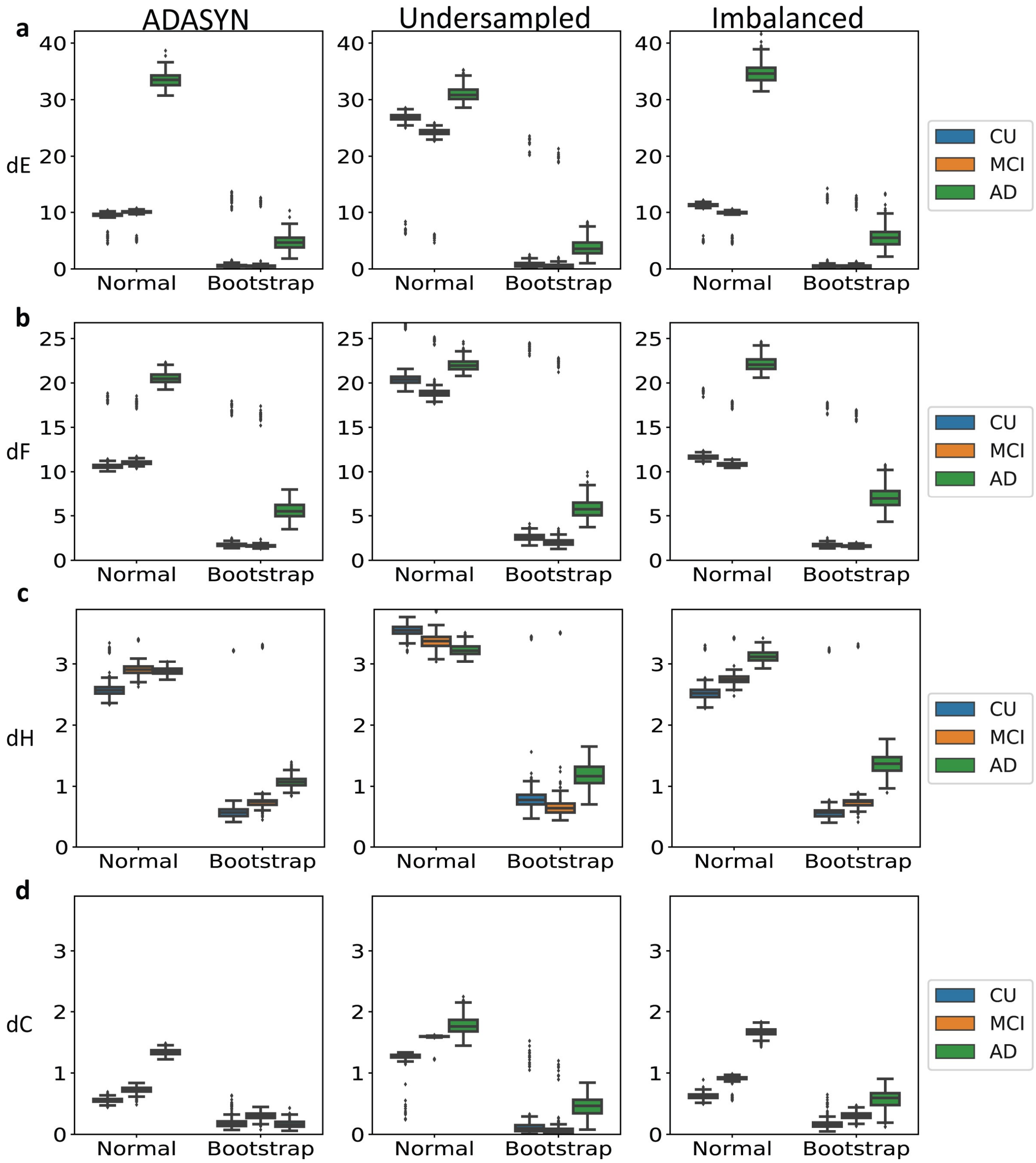


**Supplemental Figure 3. MBNs stability comparison of conventional and the MS bootstrap method for different balance schemes.** Groups CU, MCI and AD were network outlier attacked 256 times (Po = 8%) with balance schemes: ADASYN (left column), Undersampled (central column) and Imbalanced (right column). MS bootstrap MBNs were constructed using the mean matrix criterion, alpha = 0.0001 and n was set to 9300. MBNs stability evaluation using the dE, dF, dH and dC, for both conventional and bootstrap methods, are shown, respectively, in (a), (b), (c) and (d).

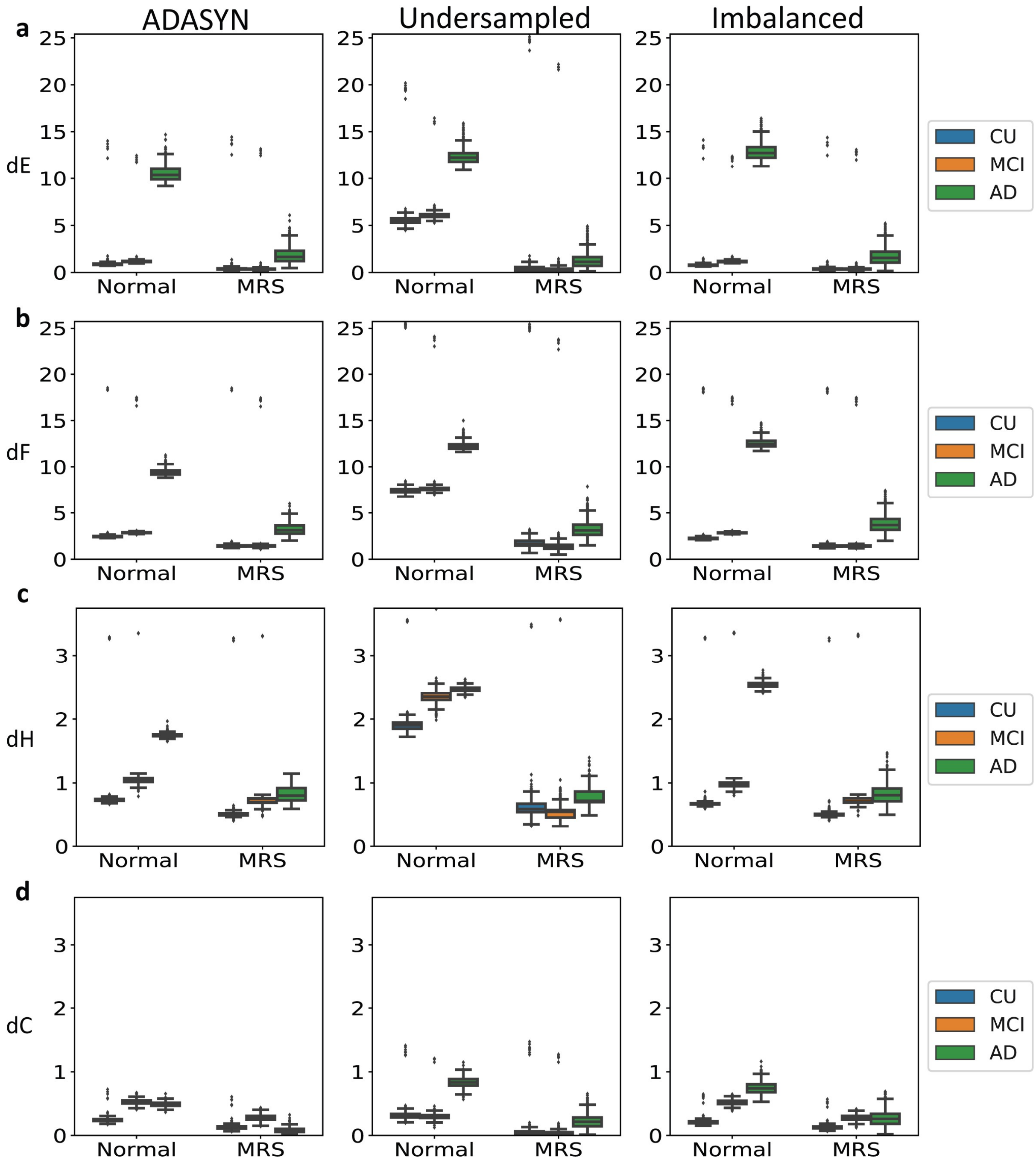


**Supplemental Figure 4. MBNs stability comparison of conventional and the MS subsampling method for different balance schemes.** Groups CU, MCI and AD were network outlier attacked 256 times (Po = 2%) with balance schemes: ADASYN (left column), Undersampled (central column) and Imbalanced (right column). MS subsampling MBNs were constructed using the mean matrix criterion, alpha = 0.0001 and n was set to 9300. MBNs stability evaluation using the dE, dF, dH and dC, for both conventional and bootstrap methods, are shown, respectively, in (a), (b), (c) and (d).

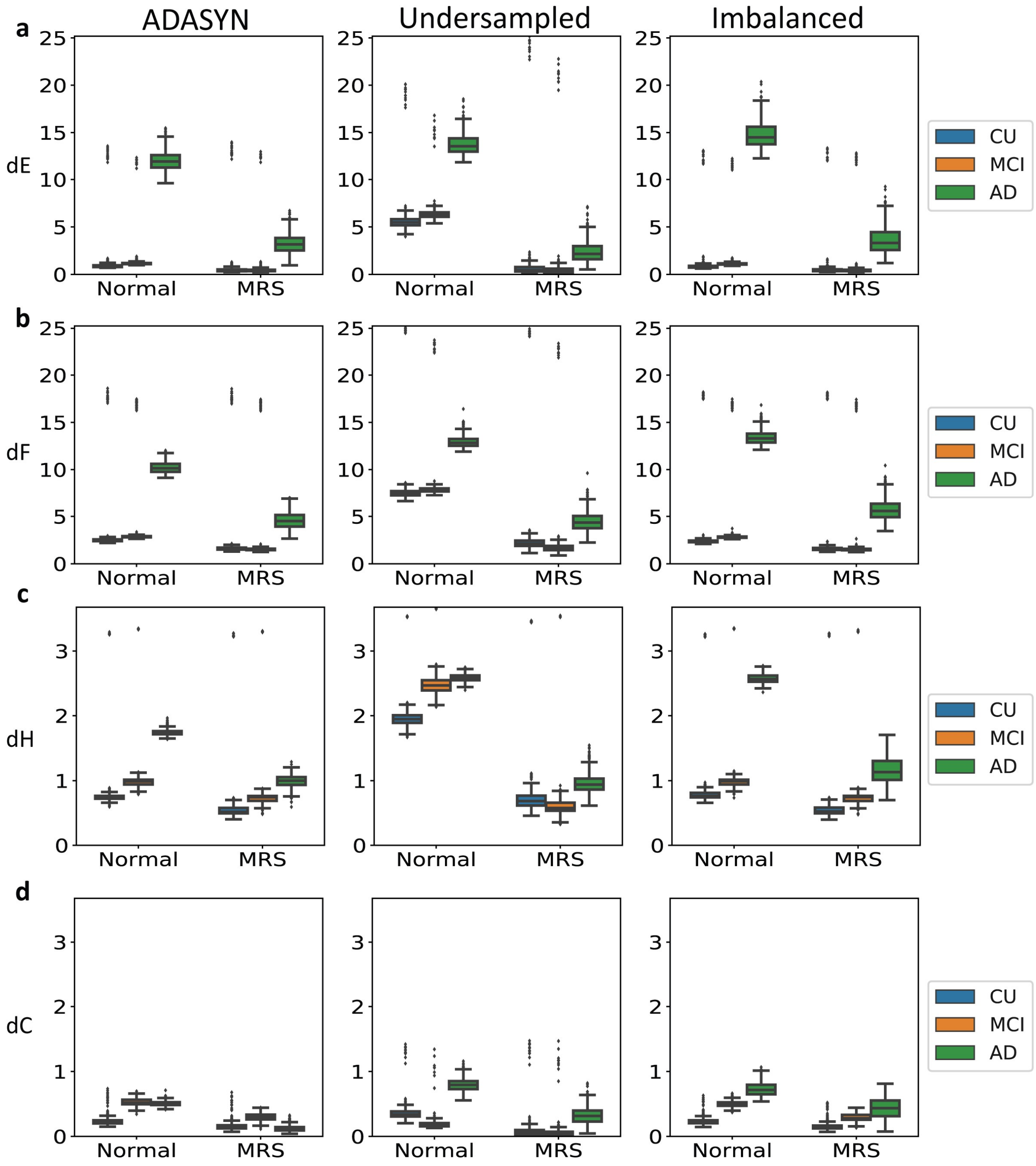


**Supplemental Figure 5. MBNs stability comparison of conventional and the MS subsampling method for different balance schemes.** Groups CU, MCI and AD were network outlier attacked 256 times (Po = 5%) with balance schemes: ADASYN (left column), Undersampled (central column) and Imbalanced (right column). MS subsampling MBNs were constructed using the mean matrix criterion, alpha = 0.0001 and n was set to 9300. MBNs stability evaluation using the dE, dF, dH and dC, for both conventional and bootstrap methods, are shown, respectively, in (a), (b), (c) and (d).

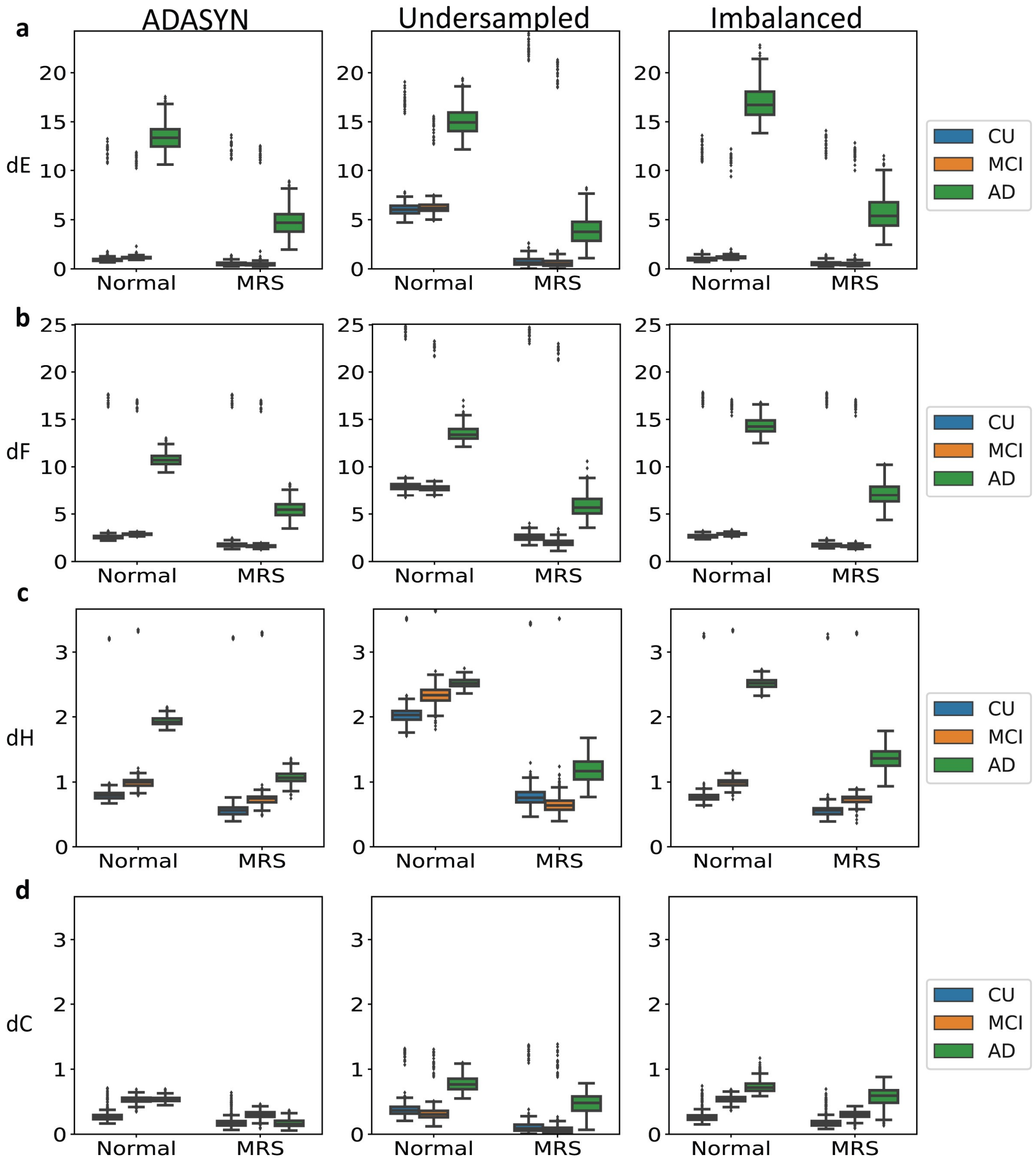


**Supplemental Figure 6. MBNs stability comparison of conventional and the MS subsampling method for different balance schemes.** Groups CU, MCI and AD were network outlier attacked 256 times (Po = 8%) with balance schemes: ADASYN (left column), Undersampled (central column) and Imbalanced (right column). MS subsampling MBNs were constructed using the mean matrix criterion, alpha = 0.0001 and n was set to 9300. MBNs stability evaluation using the dE, dF, dH and dC, for both conventional and bootstrap methods, are shown, respectively, in (a), (b), (c) and (d).

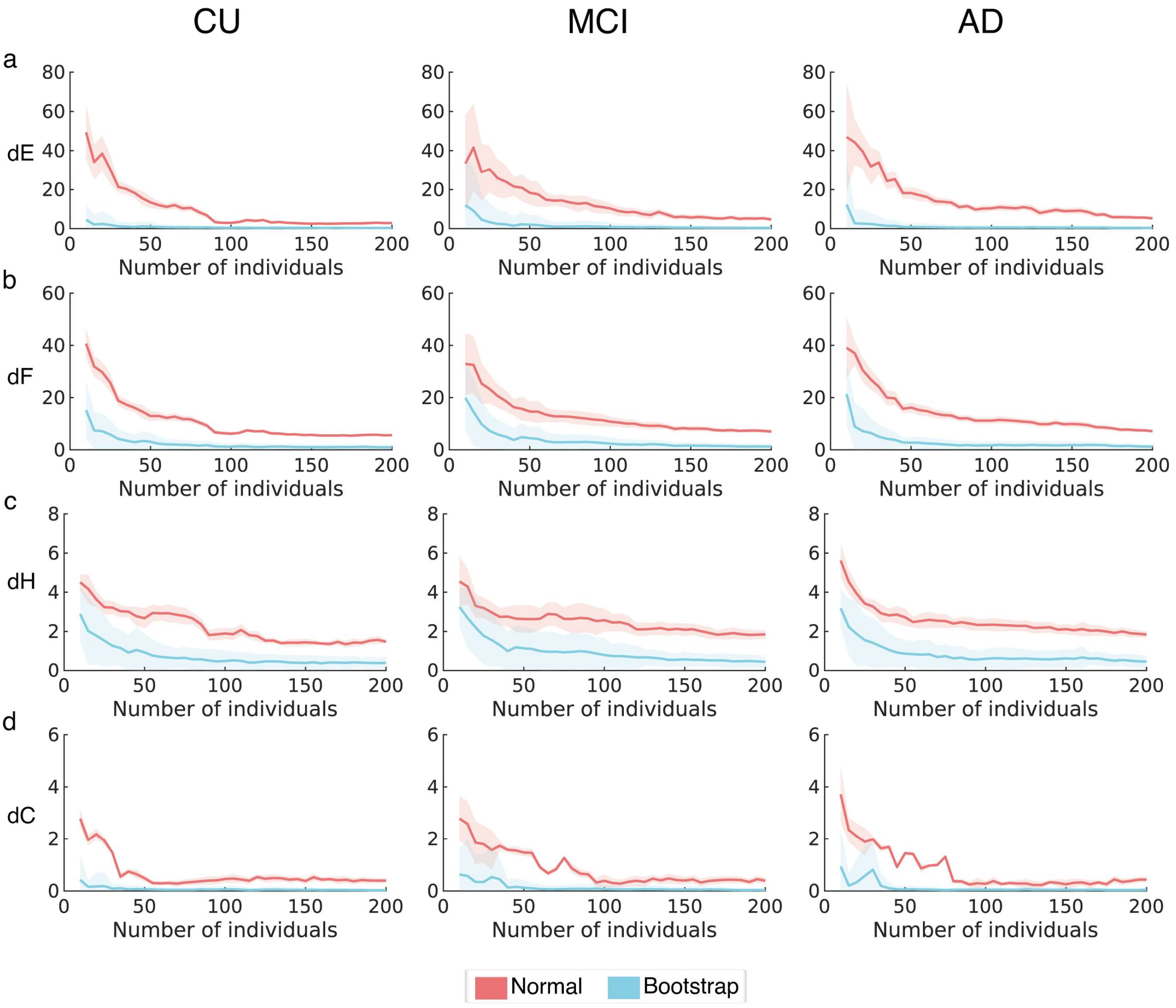


**Supplemental Figure 7. MBNs stability measures comparison of conventional and the MS bootstrap method as a function of the dataset size (i.e. number of individuals).** Groups CU, MCI and AD were network outlier attacked 256 times (Po = 5%) for each dataset size defined in the interval [10, 15, 20, …, 200]. MS bootstrap MBNs were constructed using the mean matrix criterion, alpha = 0.05 and n was set to 9300. MBNs stability evaluation using the dE, dF, dH and dC as a function of the number of individuals, for both conventional and bootstrap methods, are shown respectively in (a), (b), (c) and (d). Bold lines correspond to mean stability measure values and plotted light shadows correspond to 3 times the standard deviation from the mean measure.

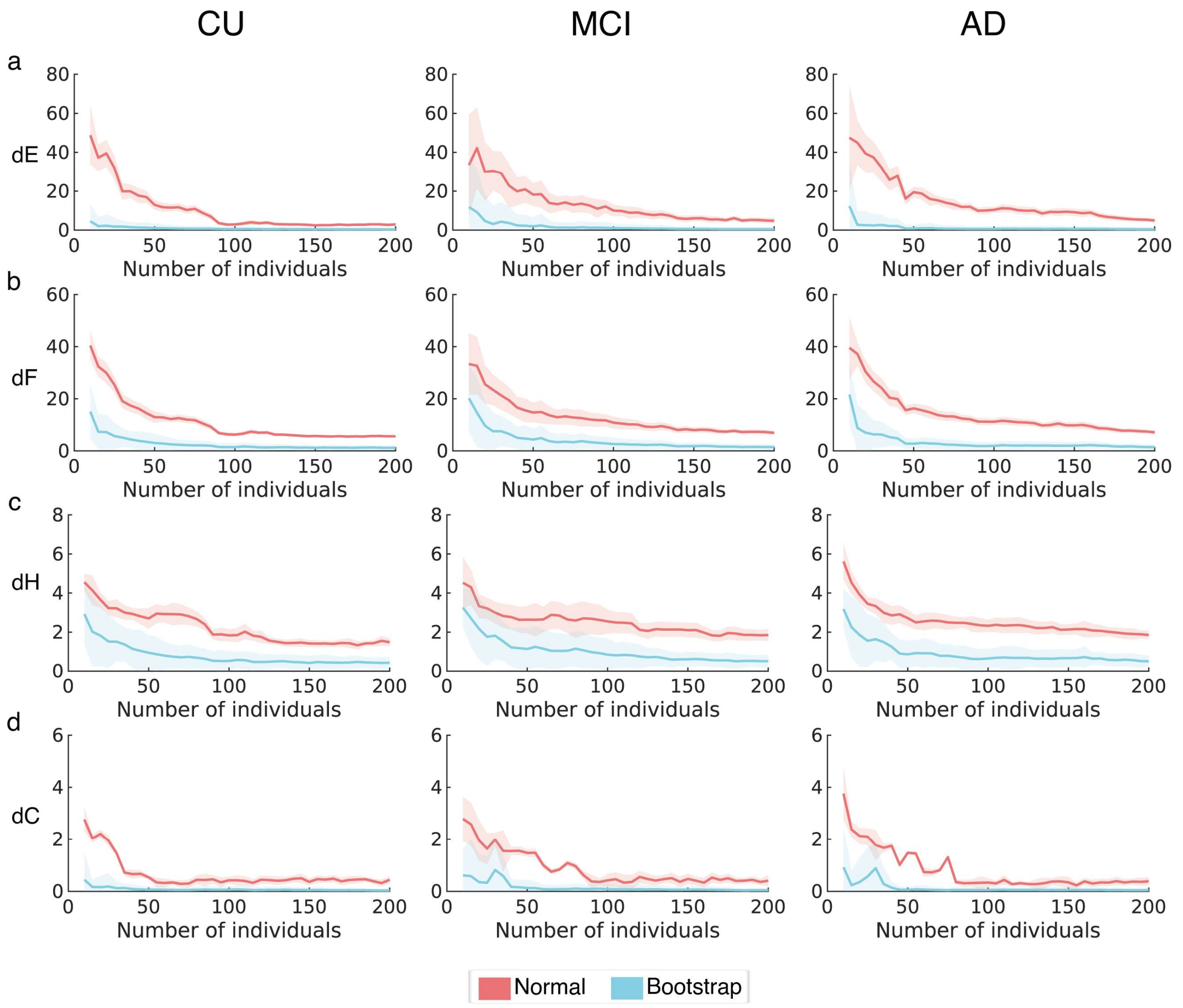


**Supplemental Figure 8. MBNs stability measures comparison of conventional and the MS bootstrap method as a function of the dataset size (i.e. number of individuals).** Groups CU, MCI and AD were network outlier attacked 256 times (Po = 8%) for each dataset size defined in the interval [10, 15, 20, …, 200]. MS bootstrap MBNs were constructed using the mean matrix criterion, alpha = 0.05 and n was set to 9300. MBNs stability evaluation using the dE, dF, dH and dC as a function of the number of individuals, for both conventional and bootstrap methods, are shown respectively in (a), (b), (c) and (d). Bold lines correspond to mean stability measure values and plotted light shadows correspond to 3 times the standard deviation from the mean measure.

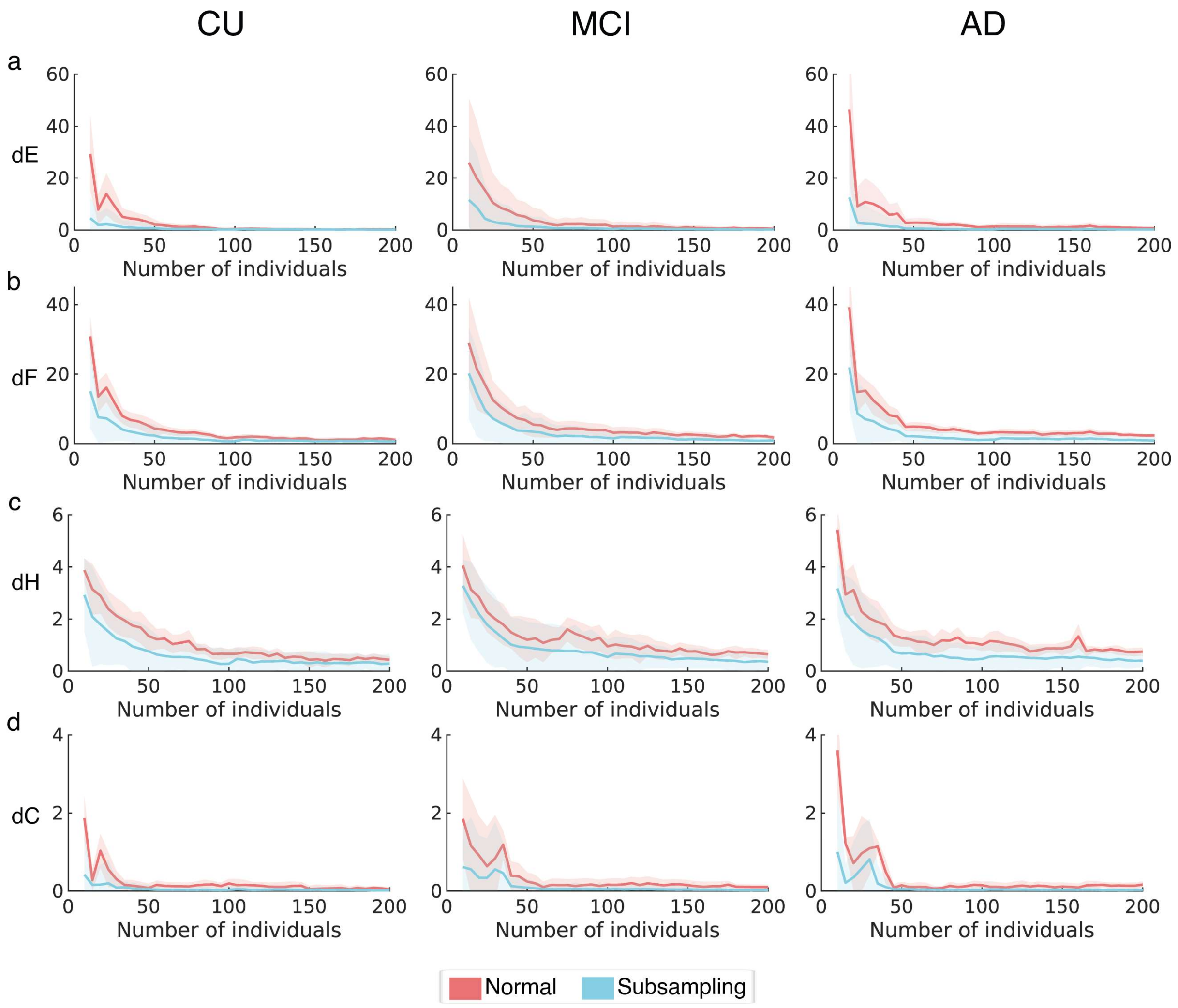


**Supplemental Figure 9. MBNs stability measures comparison of conventional and the MS subsampling method as a function of the dataset size (i.e. number of individuals).** Groups CU, MCI and AD were network outlier attacked 256 times (Po = 2%) for each dataset size defined in the interval [10, 15, 20, …, 200]. MS subsampling MBNs were constructed using the mean matrix criterion, alpha = 0.05 and n was set to 9300. MBNs stability evaluation using the dE, dF, dH and dC as a function of the number of individuals, for both conventional and bootstrap methods, are shown respectively in (a), (b), (c) and (d). Bold lines correspond to mean stability measure values and plotted light shadows correspond to 3 times the standard deviation from the mean measure.

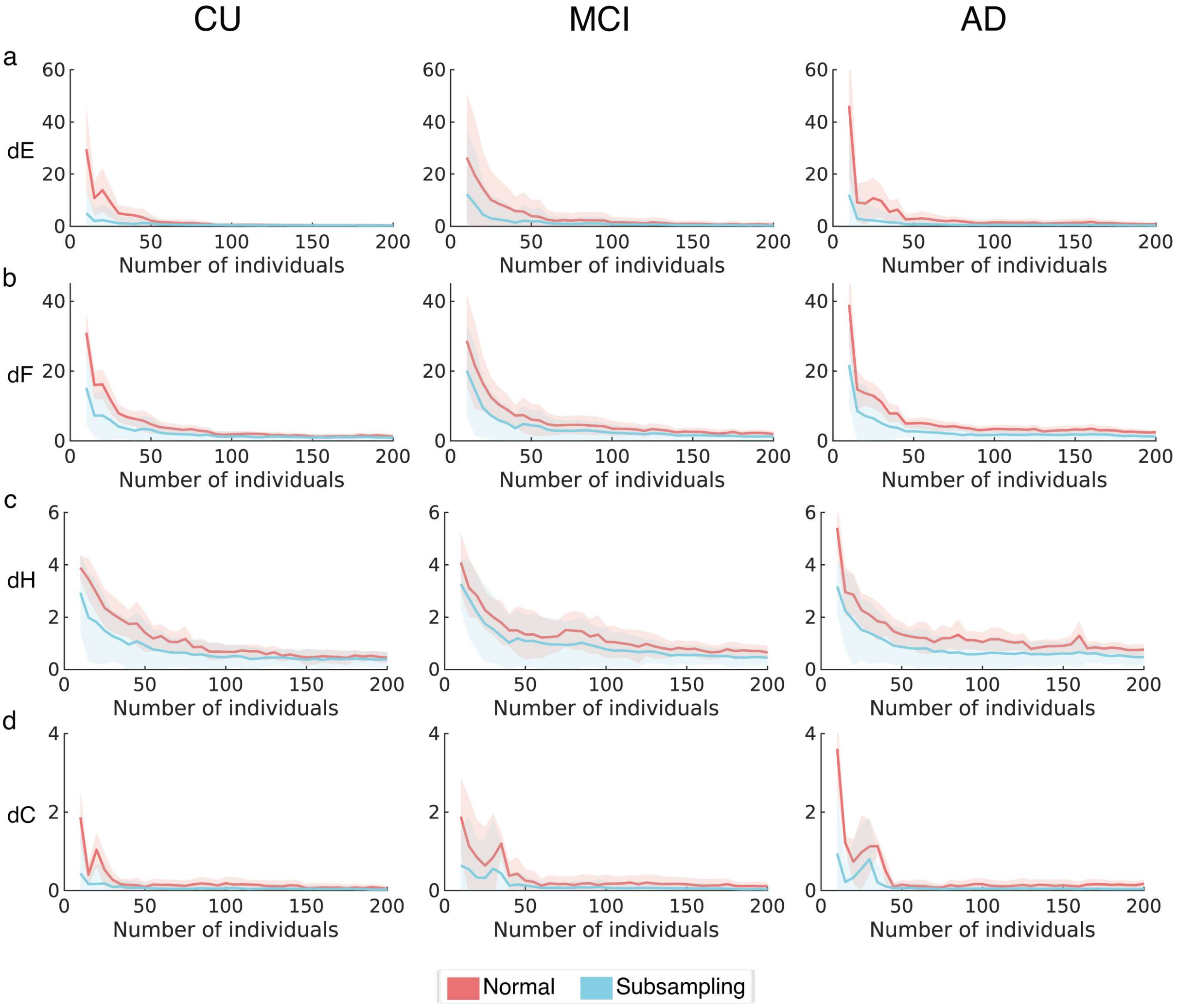


**Supplemental Figure 10. MBNs stability measures comparison of conventional and the MS subsampling method as a function of the dataset size (i.e. number of individuals).** Groups CU, MCI and AD were network outlier attacked 256 times (Po = 5%) for each dataset size defined in the interval [10, 15, 20, …, 200]. MS subsampling MBNs were constructed using the mean matrix criterion, alpha = 0.05 and n was set to 9300. MBNs stability evaluation using the dE, dF, dH and dC as a function of the number of individuals, for both conventional and bootstrap methods, are shown respectively in (a), (b), (c) and (d). Bold lines correspond to mean stability measure values and plotted light shadows correspond to 3 times the standard deviation from the mean measure.

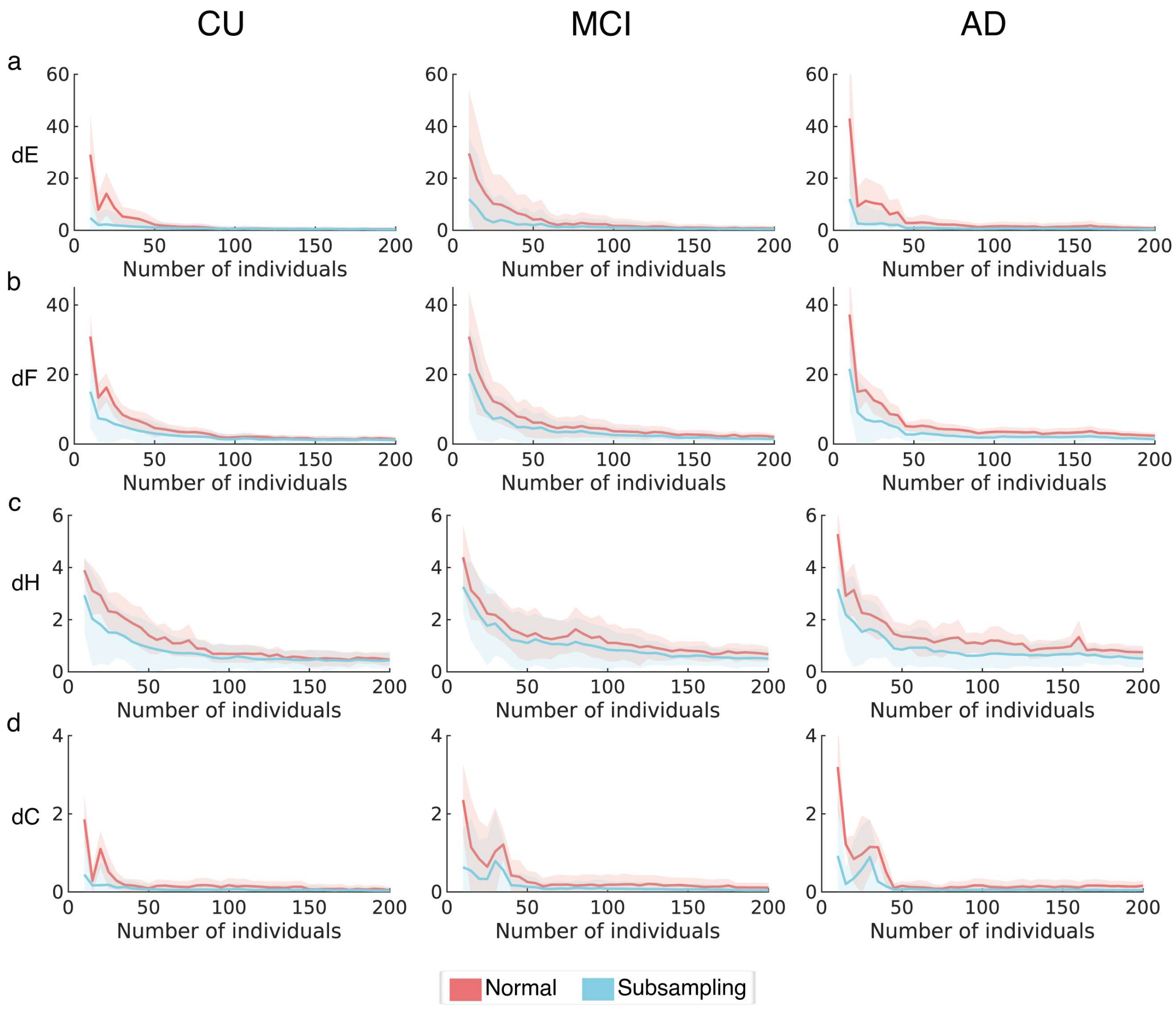


**Supplemental Figure 11. MBNs stability measures comparison of conventional and the MS subsampling method as a function of the dataset size (i.e. number of individuals).** Groups CU, MCI and AD were network outlier attacked 256 times (Po = 8%) for each dataset size defined in the interval [10, 15, 20, …, 200]. MS subsampling MBNs were constructed using the mean matrix criterion, alpha = 0.05 and n was set to 9300. MBNs stability evaluation using the dE, dF, dH and dC as a function of the number of individuals, for both conventional and bootstrap methods, are shown respectively in (a), (b), (c) and (d). Bold lines correspond to mean stability measure values and plotted light shadows correspond to 3 times the standard deviation from the mean measure.

**Table S1.** Volumes of interest.

| Adopted abbreviation | | Brain region |
|---|---|---|
| SFG_R | SFG_L | *Right and left "superior frontal gyrus"* |
| MFG_R | MFG_L | *Right and left "middle frontal gyrus"* |
| InfFG_R | InfFG_L | *Right and left "inferior frontal gyrus"* |
| MedFG_R | MedFG_L | *Right and left "medial frontal gyrus"* |
| MFOG_R | MFOG_L | *Right and left "medial front-orbital gyrus"* |
| LatFOG_R | LatFOG_L | *Right and left "lateral front-orbital gyrus"* |
| PreCG_R | PreCG_L | *Right and left "precentral gyrus"* |
| CR_R | CR_L | *Right and left "cingulate region"* |
| PostCG_R | PostCG_L | *Right and left "postcentral gyrus"* |
| SupPL_R | SupPL_L | *Right and left "superior parietal lobule"* |
| SupMg_R | SupMg_L | *Right and left "supramarginal gyrus"* |
| AngG_R | AngG_L | *Right and left "angular gyrus"* |
| Precuneus_R | Precuneus_L | *Right and left "precuneus"* |
| Ins_R | Ins_L | *Right and left "insula"* |
| SupTG_R | SupTG_L | *Right and left "superior temporal gyrus"* |
| MTG_R | MTG_L | *Right and left "middle temporal gyrus"* |
| InfTG_R | InfTG_L | *Right and left "inferior temporal gyrus"* |
| MedOTG_R | MedOTG_L | *Right and left "medial occipitotemporal gyrus/fusiform"* |
| LatOTG_R | LatOTG_L | *Right and left "lateral occipitotemporal gyrus/fusiform"* |
| ParahipG_R | ParahipG_L | *Right and left "parahippocampal gyrus"* |
| Uncus_R | Uncus_L | *Right and left "uncus"* |
| SupOG_R | SupOG_L | *Right and left "superior occipital gyrus"* |
| MOG_R | MOG_L | *Right and left "middle occipital gyrus"* |
| InfOG_R | InfOG_L | *Right and left "inferior occipital gyrus"* |
| Cuneus_R | Cuneus_L | *Right and left "cuneus"* |
| LingG_R | LingG_L | *Right and left "lingual gyrus"* |

| | | |
|---|---|---|
| OP_R | OP_L | *Right and left "occipital pole"* |
| Cerebellum_R | Cerebellum_L | *Right and left "cerebellum"* |
| HipForm_R | HipForm_L | *Right and left "hippocampal formation"* |
| Amyg_R | Amyg_L | *Right and left "amygdala"* |
| Thal_R | Thal_L | *Right and left "thalamus"* |
| SubThal_R | SubThal_L | *Right and left "subthalamic nucleus"* |
| P_R | P_L | *Right and left "putamen"* |
| Gpal_R | Gpal_L | *Right and left "globus palladus"* |
| CN_R | CN_L | *Right and left "caudate nucleus"* |
| Nacc_R | Nacc_L | *Right and left "nucleus accumbens"* |

**Table S2.** Two-way ANOVA results – MBN stability evaluation with different data imbalance setups

| **ADASYN** | | | | |
|---|---|---|---|---|
| **Effect** | **Stability Meaure** | | | |
| | **dE** | **dH** | **dF** | **dC** |
| **Group** | F (2, 1530) = 17223, p < 0.0001 | F (2, 1530) = 329.0, p < 0.0001 | F (2, 1530) = 3103, p < 0.0001 | F (2, 1530) = 8043, p < 0.0001 |
| **Method** | F (1, 1530) = 72152, p < 0.0001 | F (1, 1530) = 31514, p < 0.0001 | F (1, 1530) = 32067, p < 0.0001 | F (1, 1530) = 126559, p < 0.0001 |
| **Interaction** | F (2, 1530) = 14068, p < 0.0001 | F (2, 1530) = 20.00, p < 0.0001 | F (2, 1530) = 1541, p < 0.0001 | F (2, 1530) = 11909, p < 0.0001 |

| **Undersampled** | | | | |
|---|---|---|---|---|
| **Effect** | **Stability Meaure** | | | |
| | **dE** | **dH** | **dF** | **dC** |
| **Group** | F (2, 1530) = 243.6, p < 0.0001 | F (2, 1530) = 99.01, p < 0.0001 | F (2, 1530) = 367.0, p < 0.0001 | F (2, 1530) = 1009, p < 0.0001 |
| **Method** | F (1, 1530) = 70006, p < 0.0001 | F (1, 1530) = 65796, p < 0.0001 | F (1, 1530) = 89860, p < 0.0001 | F (1, 1530) = 59375, p < 0.0001 |
| **Interaction** | F (2, 1530) = 117.2, p < 0.0001 | F (2, 1530) = 447.6, p < 0.0001 | F (2, 1530) = 11.96, p < 0.0001 | F (2, 1530) = 744.1, p < 0.0001 |

| **Imbalanced** | | | | |
|---|---|---|---|---|
| **Effect** | **Stability Meaure** | | | |
| | **dE** | **dH** | **dF** | **dC** |
| **Group** | F (2, 1530) = 16141, p < 0.0001 | F (2, 1530) = 949.7, p < 0.0001 | F (2, 1530) = 3561, p < 0.0001 | F (2, 1530) = 6811, p < 0.0001 |
| **Method** | F (1, 1530) = 81220, p < 0.0001 | F (1, 1530) = 69351, p < 0.0001 | F (1, 1530) = 29194, p < 0.0001 | F (1, 1530) = 48148, p < 0.0001 |
| **Interaction** | F (2, 1530) = 12525, p < 0.0001 | F (2, 1530) = 66.77, p < 0.0001 | F (2, 1530) = 1550, p < 0.0001 | F (2, 1530) = 4306, p < 0.0001 |

Group factor with 3 levels: CU, MCI, AD

Method factor with 2 levels: Conventional, Bootstrap
